# Maternal group 2 innate lymphoid cells contribute to fetal growth and protection from endotoxin-induced abortion in mice

**DOI:** 10.1101/348755

**Authors:** Elisa Balmas, Batika MJ Rana, Russell S Hamilton, Norman Shreeve, Jens Kieckbusch, Irving Aye, Delia A Hawkes, Sophie Trotter, Jorge López-Tello, Hannah EJ Yong, Salvatore Valenti, Amanda N Sferruzi-Perri, Francesca Gaccioli, Andrew NJ McKenzie, Francesco Colucci

**Affiliations:** Department of Obstetrics and Gynaecology, University of Cambridge University of Cambridge School of Clinical Medicine, NIHR Cambridge Biomedical Research Centre, Cambridge CB2 0QH, UK; Department of Physiology, Development and Neuroscience, University of Cambridge, Cambridge CB2 0QH, UK; Centre for Trophoblast Research, University of Cambridge, Cambridge CB2 0QH, UK; MRC Laboratory of Molecular Biology, Francis Crick Avenue, Cambridge Biomedical Campus, Cambridge CB2 0QH, UK

**Author notes:** these authors have contributed equally (Elisa Balmas, Batika MJ Rana, Andrew NJ McKenzie, Francesco Colucci). Corresponding author: Francesco Colucci.

## Abstract

Group 2 innate lymphoid cells (ILC2s) adapt to tissue physiology and contribute to immunity, inflammatory pathology and metabolism. We show that mouse uterine ILC2s have a heightened type-2 gene signature and expand during pregnancy. Indeed, maternal ILC2s promote fetal growth and protect against fetal mortality upon systemic endotoxin challenge. Absence of ILC2s leads to utero-placental abnormalities, including poor vascular remodelling, increased *Il1b* and decreased *Il4, Il5*, and *Il13* gene expression, and reduced alternative activation of dendritic cells (DCs) and macrophages. Placentas exhibit signs of adaptation to stress, including larger maternal blood spaces and increased expression of nutrient transporter genes. Endotoxin induces the expansion of IL-1β-producing uterine DCs and, in response, more uterine ILC2s produce IL-4, IL-5 and IL-13. In a protective feedback mechanism, these cytokines suppress IL-1β-producing DCs, in line with a protective role of uILC2s against endotoxin-induced abortion. Uterine ILC2s emerge as pivotal for both normal and complicated pregnancies.

## Introduction

Poor fetal growth and infections are among the main causes of perinatal morbidity that account for 10% of the global burden of disease (WHO | The Global Burden of Disease: 2004 Update, 2014). Maternal immune cells, including lymphocytes resident in the uterus regulate fetal growth and participate in defense against pathogens, however it is unclear how they achieve the right balance between trophic and immune functions (Shmelova & Colucci, 2021; Erlebacher, 2013). Both heterogeneity and tissue specialization of immune cells may hold the key to understanding the pathophysiology of some important pregnancy complications, such as miscarriage and fetal growth restriction (FGR), which is defined as the failure of the fetus to achieve its genetically determined growth potential. Consequently, it is critical that we understand the cellular and molecular immune factors that regulate the uterine environment. Innate lymphoid cells (ILCs) lack antigen receptors but are equipped with receptors for cytokines, hormones, and paracrine signals enabling them to sense their environment and respond quickly by producing factors that influence parenchymal cells and other resident immune cells. Despite often comprising only a small proportion of tissue resident immune cells, ILCs are present in most tissues and have been shown to orchestrate inflammatory responses to viruses, intracellular bacteria and parasites. Through alternative activation of macrophages and other myeloid cells, ILCs also drive and support repair mechanisms that maintain tissue integrity (Artis & Spits, 2015; Kim & Van Dyken, 2020; Xiong et al., 2022).

The diverse responses of ILCs can be ascribed to three main groups of cell populations defined by their differential expression of the transcription factors and cytokines they respond to and produce (Eberl et al., 2015). Group 2 ILCs (ILC2s), are characterized by the transcription factor GATA-3 and the IL-33 specific receptor ST-2. They respond to IL-25, IL-33 and TSLP and produce both amphiregulin, which is involved in tissue repair, and classical type-2 cytokines including IL-4, IL-5 and IL-13. Through the production of these soluble factors and by interacting with other immune cells, such as eosinophils, type-2 helper T cells and alternatively activated macrophages and dendritic cells (DCs), ILC2s are attuned to the physiology of the tissue in which they differentiate (Jenkins et al., 2013; Kim & Van Dyken, 2020; Molofsky et al., 2013; Nussbaum et al., 2013; Spitz & Mjosberg, 2022; Xiong et al., 2022).

We, and others, have described ILCs resident in the uterus before and during pregnancy, in human and mice (Doisne et al., 2015; Miller et al., 2021; Sugahara et al., 2022; Vacca et al., 2014; Xu et al., 2018). While functions have been suggested for uterine group 1 and 3 ILCs in mice or humans (Ashkar et al., 2000; Boulenouar et al., 2016; Croxatto et al., 2016; Montaldo et al., 2015), the physiological roles of ILC2s in normal and complicated pregnancies remain poorly understood, also because very few if any ILC2s are found in human endometrium or decidua in early pregnancy (Doisne et al., 2015; Vento-Tormo et al., 2018; Huhn et al., 2020). *Nfil3*^*-/-*^dams completely but not selectively lack ILC2s and have impeded vascular remodelling of the uterine arteries, inefficient placentas, and growth restricted fetuses (Boulenouar et al., 2016). Uterine ILC2s respond to the alarmin IL-33 and are regulated by female sex hormones in mice (Bartemes et al., 2017; Miller et al., 2022). Pups from dams lacking the IL-33 receptor ST-2 have increased neonatal mortality (Bartemes et al., 2017). Interestingly, polymorphisms in genes coding for cytokines of the IL-1 family, including *IL1B* and *IL33*, are associated with recurrent miscarriage in certain populations(Yue et al., 2016; Zhang et al., 2017) and ILC2 are increased in the decidua of women with preterm labour (Xu et al., 2018). Both IL-33 and ST-2 have been implicated in pre-eclampsia (Granne et al., 2011). Moreover, IL-33 and maternal lymphocytes are linked to preterm labour in both humans and mice (Huang et al., 2016). Thus, there is emerging evidence that IL-33 and ILC2s are involved in both healthy and complicated pregnancies but more investigation is required to understand their role in reproduction (Miller et al., 2021; Miller et al., 2022).

Dysregulation of cytokines and immune cell types in the uterus may impair placental function, thus triggering pregnancy syndromes, including FGR and miscarriage. A number of fetal abnormalities can be ascribed to malfunctioning placentas(Perez-Garcia et al., 2018) and the maternal environment influences the interdependent growth of both fetus and placenta (Sferruzzi-Perri & Camm, 2016). Indeed, a reduction in the ratio between fetal and placental weight generally reflects reduced placental efficiency (Fowden et al., 2009). Placental insufficiency is linked to neurodevelopmental disorders(Rehn et al., 2004). In fetuses that fail to attain their genetically determined growth potential, a circulatory compensation occurs where changes in cerebral blood flow allow for the brain to develop despite FGR. This seemingly advantageous ‘brain sparing effect’ is however linked to cognitive defects later in life (Scherjon et al., 2000). Maternal immune activation also influences fetal brain development(Smith et al., 2007) and dysregulation of uterine immune cells, such as ILC2s, may adversely affect fetal development of the brain.

Based on the evidence that IL-33 and ST-2 are involved in pregnancy complications (Miller et al., 2021; Miller et al., 2022) and considering the prominent roles of ILC2s in homeostasis and immunity in lung, intestine, and adipose tissue (Klose & Artis, 2016), we hypothesized that uterine ILC2s play essential roles in normal and complicated pregnancies. Our data show that ILC2s have unique features in the mouse uterus. Whole genome transcriptome analysis shows that the typical type-2 immunity gene signature characterized by GATA-3, ST-2 and IL-5 is enhanced in uterine ILC2s compared to lung and lymph nodes. Using a mouse model of selective ILC2 deficiency, we show that the uterine immune environment of ILC2-deficient dams is dysregulated and pro-inflammatory. Pregnancies in ILC2-deficient dams are characterized by growth-restricted fetuses. Moreover, due to the unbalanced uterine immunological environment, ILC2-deficient dams are more susceptible to fetal demise when exposed systemically to bacterial endotoxin. Together, our data demonstrate uILC2s as master regulators of type-2 immunity in the mouse uterus and as key immune cells in the maintenance of immunological homeostasis during both normal and complicated pregnancies.

## Results

### ILC2s colocalize with IL-33 in the mouse uterus and are found in the human uterus

The mouse uterus contains ILC2s (uILC2s) that reside in the muscle layer of the uterine wall (myometrium), and expand over 5-fold, from about 500 to 3,000 cells, during pregnancy (**Fig. 1A** and **S1A**). Since IL-33 is a known driver of ILC2 expansion, and GWAS studies have linked *IL33* to pregnancy abnormalities, we examined *Il33*-citrine reporter mice (*Il33*^*cit/wt*^) for the presence of IL-33 expression in the uterus (Hardman et al., 2013). Indeed, we identified *Il33*-citrine^+^ cells, mainly localized in the myometrium, where ILC2s reside, with some *Il33-*citrine expression also in the decidua (**Fig. 1B**). Cells expressing *Il33* were restricted to the stromal CD45^-ve^ population in the myometrium at both midgestation (gestation day 9.5 = gd9.5) and term (gd18.5) (**Fig. S1B, C**), and this is consistent with IL-33 protein levels in the uterus (**Fig. S1D**). Dams deficient in both copies of *Il33* (*Il33*^*cit/cit*^, hereafter referred to as IL-33KO mice) have a 90% reduction in the total number of uILC2s (**Fig 1C**), demonstrating the vital role that IL-33 plays in ILC2 homeostasis in the uterus. Although similar to the mouse, human CD127^hi^ uILC2s are virtually absent in the uterine mucosa (Doisne et al 2015), we have now identified uILC2s within CD127^low/-ve^ cells in both the non-pregnant endometrium and the decidua (**Fig 1SE, F**). These results show that both murine and human uterine tissues contain small populations of uILC2s, which, like in other tissues, in the mouse depend on and colocalise with IL-33.

**Figure 1.**
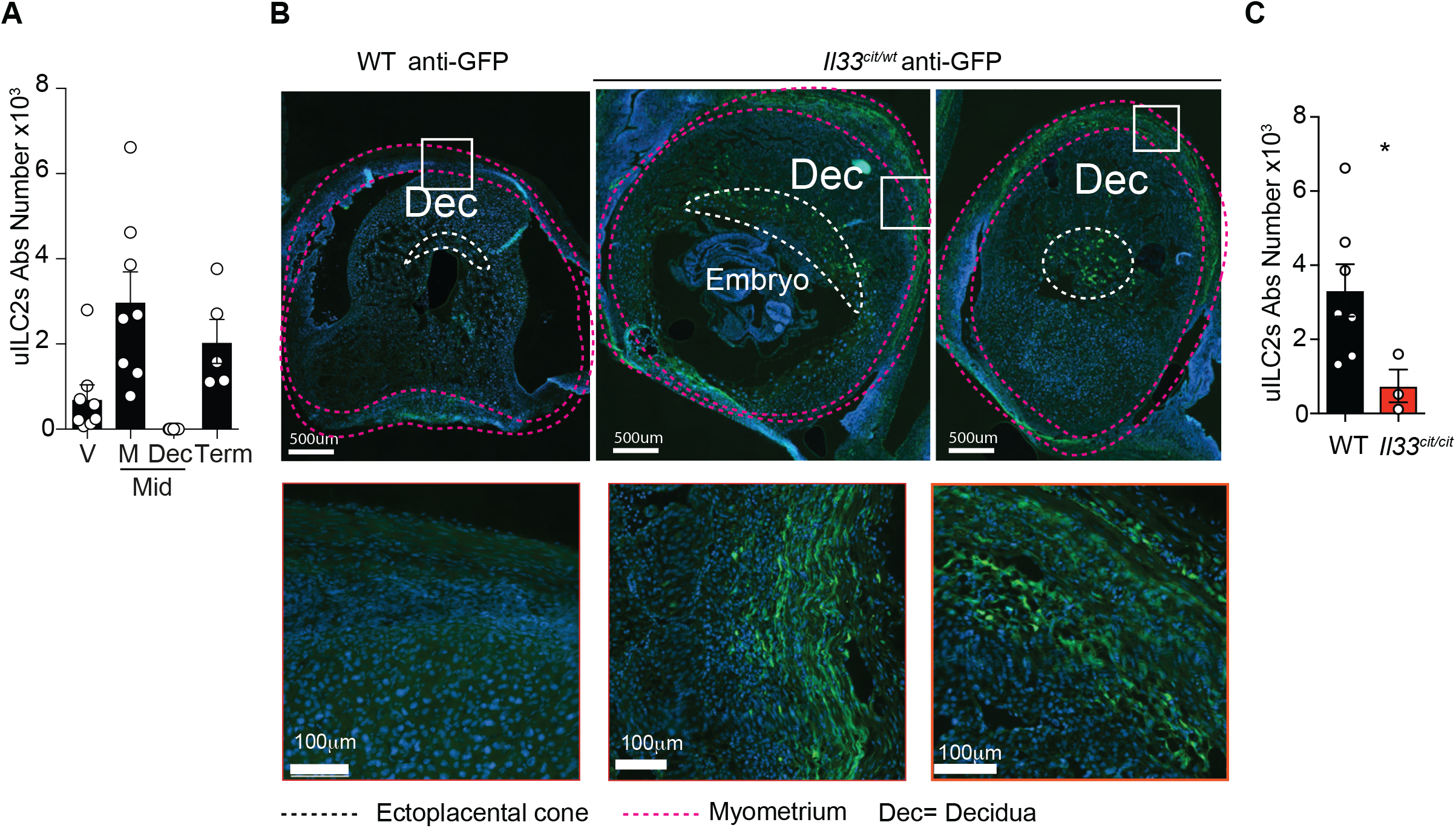
ILC2s co-localise with IL-33 in the mouse uterus. **(A)** Absolute numbers of uterine ILC2s determined by flow cytometry <u>in virgin (V) mice or dams from ♀WT x ♂WT crosses at mid-gestation</u> (Mid = gestation day 9.5 = gd9.5), and at Term (gestation day 18.5 = gd18.5) (n=5-8). **(B)** Immunohistochemistry of implantation sites in dams from ♀*Il33*^*cit/wt*^ x *Il33*^*cit/wt*^ ♂ crosses <u>at mid-gestation</u>. Myometrium is indicated in red, and the ectoplacental cone in white; **(C);** Absolute numbers of uterine ILC2s determined by flow cytometry in dams from ♀WT x ♂WT crosses (n=7) and dams from ♀*Il33*^*citcitt*^ x *Il33*^*cit/wt*^ ♂ crosses (n=3 pools of 5 uteri/replicate) <u>at mid-gestation</u>. Data in A and C are displayed as mean ± SEM. Data in A were analysed with Kruskal-Wallis test and data in C were analysed by Student’s *t-test*.

### Fetal growth restriction in fetuses of ILC2-deficient dams

Since uILC2s expand during pregnancy and ILC2s are known regulators of tissue physiology, we reasoned that, despite the small number of uILC2 in the uterus, they may also play important roles in reproduction and fetal development. To test this hypothesis, we utilised *Rora*^*flox/flox*^*Il7ra*^*cre/wt*^ ILC2-deficient mice (hereafter referred to as ILC2KO mice), in which the RORα transcription factor that is critical for ILC2 development is deleted in IL7Rα-expressing ILC2s (Oliphant et al., 2014; Schlenner et al., 2010). As expected, maternal uILC2s were undetectable in the uterus of ILC2KO females mated with wild-type males (**♀**ILC2KO x ♂WT), whereas normal numbers of maternal uILC2s at mid-gestation were present in the uterus of WT females mated with ILC2KO males (**♀**WT x ♂ILC2KO) (**Fig. 2A**). It is important to note that the genetic make-up of the fetuses and their placentas are equivalent in both crosses. Strikingly, although gestation goes to term with unaltered litter sizes and low perinatal mortality in both **♀**ILC2KO x WT and reverse **♀**WT x ♂ ILC2KO cross (Fig. **S2A**), fetuses at term (gd 18.5) from **♀**ILC2KO x ♂WT cross are on average about 8% lighter than fetuses from the reverse **♀**WT x ♂ILC2KO cross (1.105+/- 0.01 gr and 1.203+/-0.008 gr, respectively, **Fig 2B**). Thus, the absence of maternal ILC2s, and not the fetal genotype, is responsible for intrauterine fetal growth restriction (FGR) in this model. Similarly, the **♀**IL-33KO x ♂WT cross produce lighter fetuses (1.089+/-0.018 gr), further supporting the hypothesis that the maternal IL-33-ILC2 pathway is important for optimal fetal growth. The ratio between fetal weight and placental weight is a gross indicator of placental efficiency (Fowden et al., 2009), with the fetus weighing normally about 15 times its placenta at term. Interestingly, we found that both the ♀ILC2KO x ♂WT and the **♀**IL-33KO x ♂WT crosses result in significantly reduced fetal/placental weight ratio compared with those from control crosses (**Fig. S2B**). Moreover, the lack of maternal IL-33 also affects placental growth as the placentas generated in the **♀**IL-33KO x ♂WT crosses are heavier than controls (**Fig. 2B**). FGR is usually caused by placental insufficiency, to which fetuses respond with the brain-sparing effect, a phenomenon that spares the fetal brain from hypoxia but exposes it to neurological disorders later in life. Notably, measurements of 3D-reconstructed CT-scan images of fetus head demonstrated that fetuses carried by dams lacking ILC2s (**♀**ILC2KO x ♂WT) display significantly smaller head diameter than those carried by ILC2 sufficient dams (**♀**WT x ♂ILC2KO, **Fig. 2C-D**). The brain volume, but not the femur length of these fetuses is also reduced (**Fig. 2C-E**), suggesting that bone development may be less affected than brain development. However, the ratio between brain volume and the total body weight is similar in fetuses from both crosses, suggestive of overall symmetric FGR (**Fig. S2C**). Taken together, these data highlight the importance of the maternal IL-33-ILC2 pathway in placental efficiency and fetal growth.

**Figure 2:**
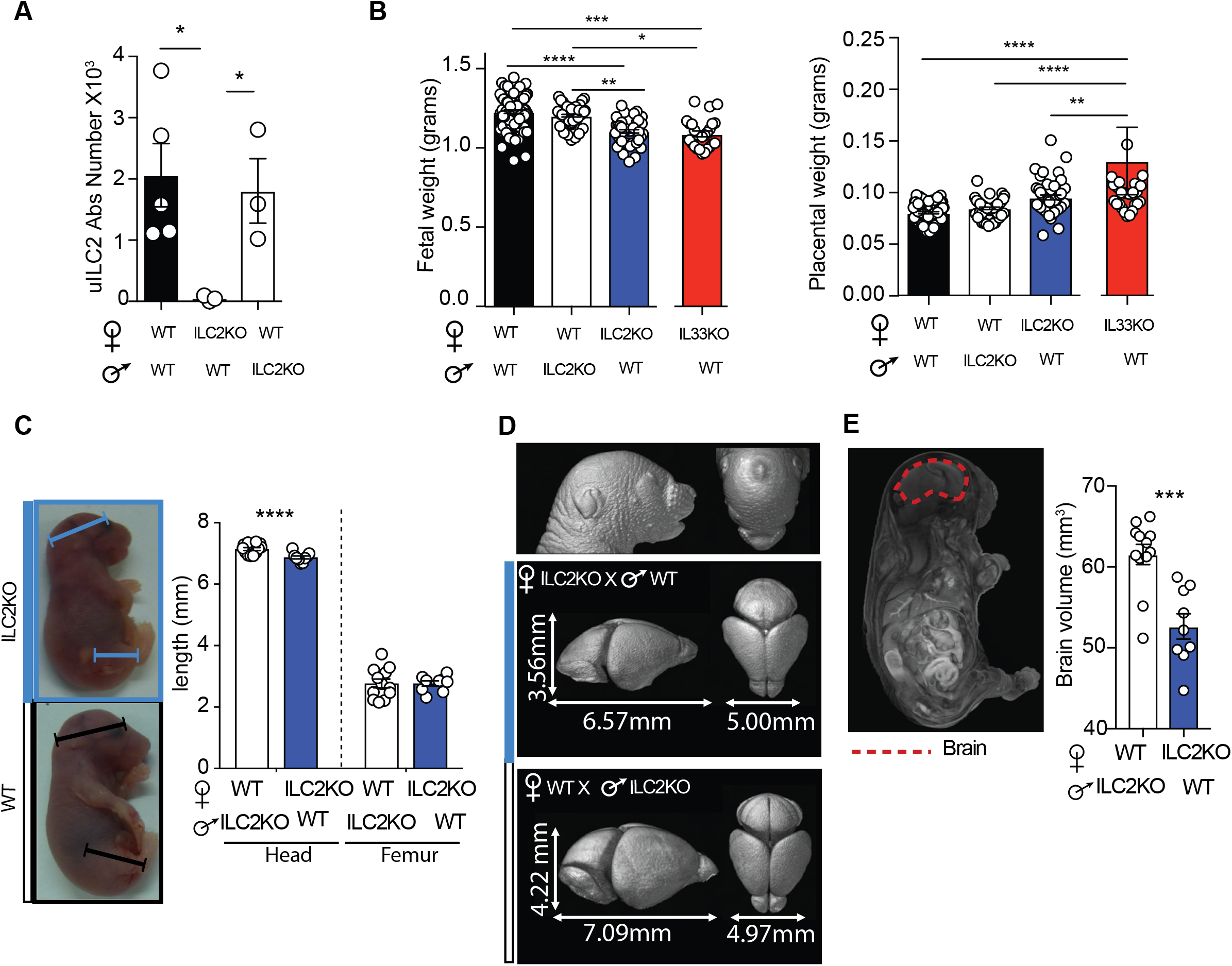
Fetal growth restriction in foetuses of ILC2 KO dams. **(A)** Flow cytometry data showing uILC2 numbers at mid-gestation in dams from ♀WT x ♂WT (n=5), ♀ILC2KO x ♂WT (n=3) and ♀WT x ♂ILC2KO (n=3) crosses. **(B)** Fetal and placental weights from ♀WT x ♂WT (n=13 litters), ♀ILC2KO x ♂WT (n=8 litters); ♀IL33KO *(Il33*^*cit/cit*^) x ♂WT crosses (n=4 litters); ♀WT x ♂ILC2KO (n=7 litters) crosses. Significance was determined using a mixed model approach that accounts for inter-litter variability but not sex. **(C)** Measurements of the head diameter and brain segmentation (the measurements are representative of one brain for cross) and femur length per cross and their quantification. **(D)** 3D Micro-CT imaging of gd18.5 fetuses from ♀ILC2KO x ♂WT crosses (n=9 from 3 dams) and ♀WT x ♂ILC2KO crosses (n=12 from 4 dams). **(E)** Example of the brain volume demarcation (red, left) and its quantification (right). Data are displayed as mean ± SEM. Data in A and B were analysed by Kruskal-Wallis test, data in D and E were analysed in each tissue by paired Student’s *t-test*.

### Abnormal utero-placental adaptations to pregnancy in ILC2KO dams

In an attempt to determine the nature of FGR in fetuses of ILC2-deficient dams, we analysed the decidua (**Fig. 3A-C**) and the placenta (**Fig. 3D and E**) of these mice to seek evidence of abnormalities in the utero-placental unit. Uterine arteries undergo a critical transformation early in human and murine gestation, so that the lumen increases to provide blood supply to the placenta. Strikingly, in ILC2KO dams the decidual vessels/lumen ratio is greater than in WT dams (**Fig. 3A**), suggesting insufficient widening of the uterine arteries. During uterine vascular remodelling, arterial smooth muscle actin becomes degraded at mid-gestation, a process readily visible in WT dams, but clearly less pronounced in ILC2KO dams (**Fig. 3B**), confirming sub-optimal vascular changes. ILC2s secrete type-2 cytokines and in their absence the uterine immune microenvironment may be dysregulated. We measured transcripts of inflammatory cytokines and found significantly enhanced expression of *Il1b* in the decidua of ILC2KO dams, suggesting a pro-inflammatory microenvironment (**Fig. 3C**). Furthermore, we found that the expression of 4 genes coding for glucose and amino acid transporters (*Glut1, Glut3, Snat2* and *Snat4*) is enhanced in term placentas of ILC2KO dams, suggesting compensatory placental mechanisms occur in an attempt to optimise the supply of nutrients in the face of an adverse environment caused by maternal ILC2 deficiency (**Fig. 3D**). Increased maternal blood spaces in the placentas of ILC2KO dams also support this idea (**Fig. 3E** and **Fig. S3, and Table S1**). These results demonstrate that maternal ILC2s contribute to the widening of the decidual vessels and also counteract the expression of the pro-inflammatory cytokine IL-1β in the decidua, two roles that can affect placental function. Taken together these data suggest the placentas adopt compensatory mechanisms which serve to optimise nutrient and oxygen delivery to the fetus in ILC2KO dams.

**Figure 3:**
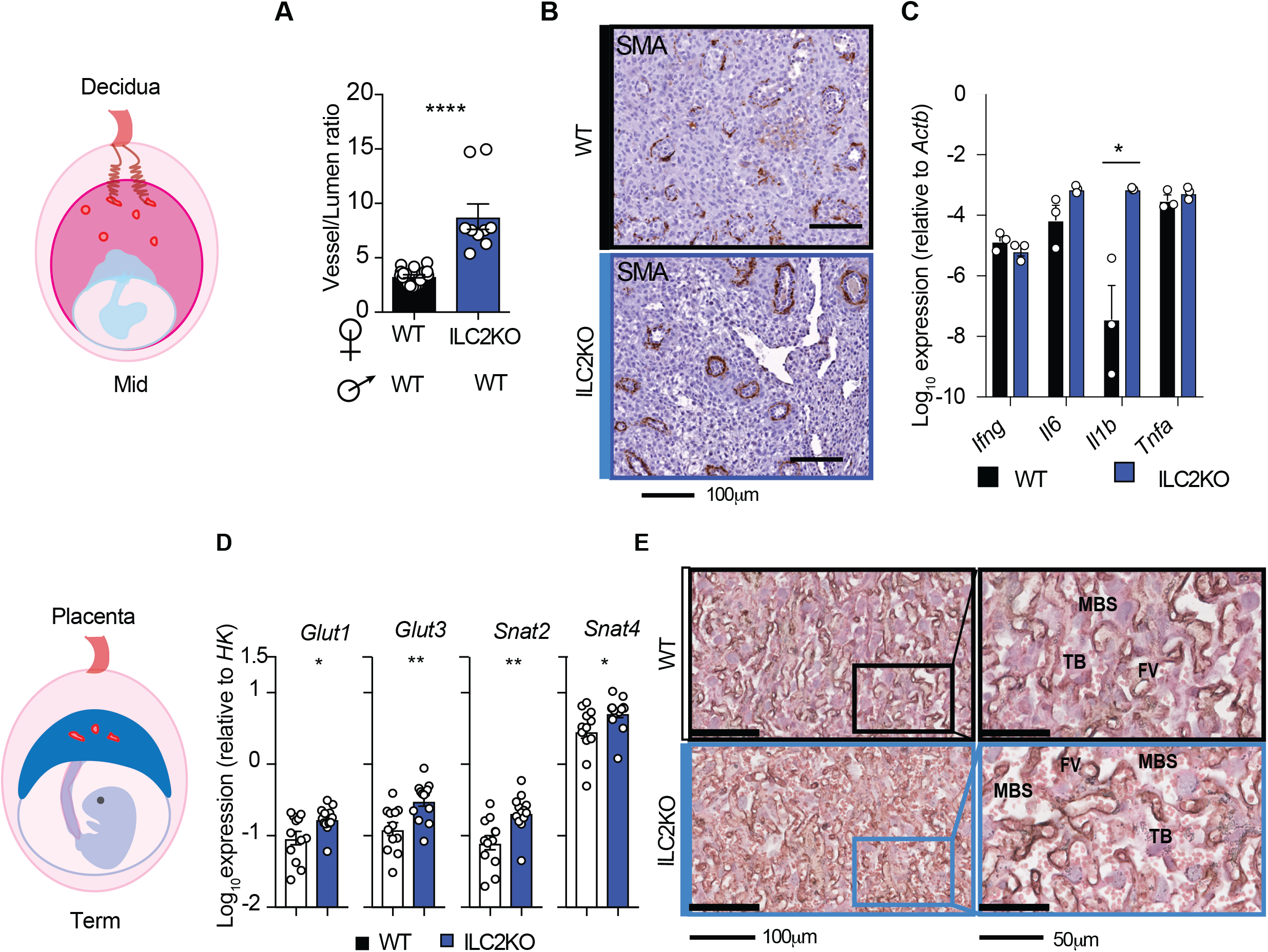
Decidual and placental abnormalities in ILC2 KO dams. **(A)** Stereological and immunohistochemical quantification of uterine artery thickness and lumen area ratios in the decidua of dams from ♀ILC2KO x ♂WT crosses and dams from ♀WT x ♂WT crosses at midgestation (gd9.5). Pooled data from 4 litters per cross, n=12–20 total implantation sites per cross. **(B)** Smooth muscle actin (SMA) staining on decidual uterine arteries at midgestation (gd9.5) in dams from ♀WT x ♂WT and ♀ILC2KO x ♂WT crosses; 12.3X magnification, scale bar 100μm. **(C)** Relative *mRNA* expression of pro-inflammatory cytokines in the uterus of dams from ♀WT x ♂WT (n=4) and ♀ILC2KO x ♂WT (n=3) crosses at midgestation (gd9.5). **(D)** Relative *mRNA* expression of tissue glucose and amino acid transporters from dissected term placentas (gd18.5). of 4 ♀WT x ♂ILC2KO crosses (n=12 placentas) and 5 ♀ILC2KO x ♂WT crosses (n=13 placentas). **(E)** Representative photomicrographs of sections of placentas at term (gd18.5) stained for lectin, cytokeratin and eosin to highlight fetal vessels (FV), trophoblast (TB) and maternal blood spaces (MBS);. Data in A, C and D are displayed as mean ± SEM. Data in A were analysed by Mann-Whitney test, data in C and D were analysed by paired Student’s *t*-test.

### Comparative transcriptome analysis of uterine, lymph node and lung ILC2s

We next turned to a molecular analysis to understand functions that may explain the contribution of uILC2s to the utero-placental unit. Considering the previously reported differences in the functional profile of ILC2s in the uterus and other tissues (Doisne et al., 2015), we set out to analyse the transcriptome of uILC2s to determine their molecular identity. To do this, we compared the genome-wide transcriptome of murine ILC2s of the uterus (from virgin WT mice and from dams of **♀**WT x ♂WT crosses, at mid-gestation and at term), the lung and the lymph nodes (mesenteric, para-aortic and inguinal lymph nodes pooled together) of virgin WT mice. Principal component analysis showed that uILC2s cluster separately from ILC2s purified from the lung and the lymph nodes (**Fig. 4A**), suggesting uterine-specific features. To identify uterine-specific genes in uILC2s we generated a gene list comparing the three uterine ILC2 populations to those of lung and lymph nodes. We found that 3,279 genes were upregulated in uILC2s (**Fig. 4B**). These included canonical ILC2-signature genes such as *Gata3* and *Il1rl1* (coding for the IL-33 receptor ST-2), as well as *Il4, Il5, Il13*, and *Areg* coding for Amphiregulin, which could regulate trophoblast growth (Lysiak et al., 1995). High levels of *Arg1* coding for Arginase-1 are also detected in uILC2s. Arginase-1 regulates bioenergetics and type-2 immune responses in lung ILC2s (Monticelli et al., 2016) (**Fig. 4B,C**). Interestingly, as well as being more highly expressed in uILC2s, these type-2 genes also increase in uILC2s as pregnancy progresses, with highest expression levels at term. Genes associated with immunoregulation are also upregulated in uILC2s, including *Ctla4, Foxp3, Tgfb1* and *Tgfb2* (**Fig. 4B,D**). Heat maps of expression of these selected genes across gestation and, in comparison with lung and lymph node, showed that the enhanced type-2 signature and the expression of immunomodulatory genes in uILC2s both increase with gestation (**Fig. 4C, D**). Protein expression analysis by flow cytometry supported RNA-seq findings that uILC2 have an enhanced type-2 signature and distinct functions compared to lung resident ILC2s. uILC2s showed higher expression of IL-5 and ST-2 in response to IL-25 and IL-33, as well as constitutively higher expression of CTLA-4 and FoxP3, than lung ILC2s (**Fig. S4A-C**). Other genes upregulated in uILC2s include non-cytotoxic granzymes *Gzma, Gzmc, Gzmd, Gzme, Gzmf, Gzmg, Gzmk*, most likely involved in tissue remodeling and *Csf1*, encoding CSF-1, a crucial regulator of macrophages and DC dynamics in the uterus (Tagliani et al., 2011). In addition, Gene Ontology analysis also highlighted the expression of genes involved in controlling recruitment and homeostasis of T cells, eosinophils and myeloid cells, such as *Ccl5, Csf1, Csf2, Csf2rb, Csf2rb2, Csf3, Cxcl2*, as well as genes mediating the response to oestrogens and estradiol and the oestrogen receptor *Ers1* (**Fig. S4D** and **Table S2**). The combined higher expression of GATA-3, IL-5 and ST-2 in uILC2s showcases their enhanced type-2 profiles compared to lung and lymph node. This suggests that uILC2s have uterine-specific regulatory functions that may support gestation. Moreover, the expression of genes likely to regulate other immune cells suggest that the role of uILC2s in gestation may also be mediated through interaction with these cells in the uterus.

**Figure 4:**
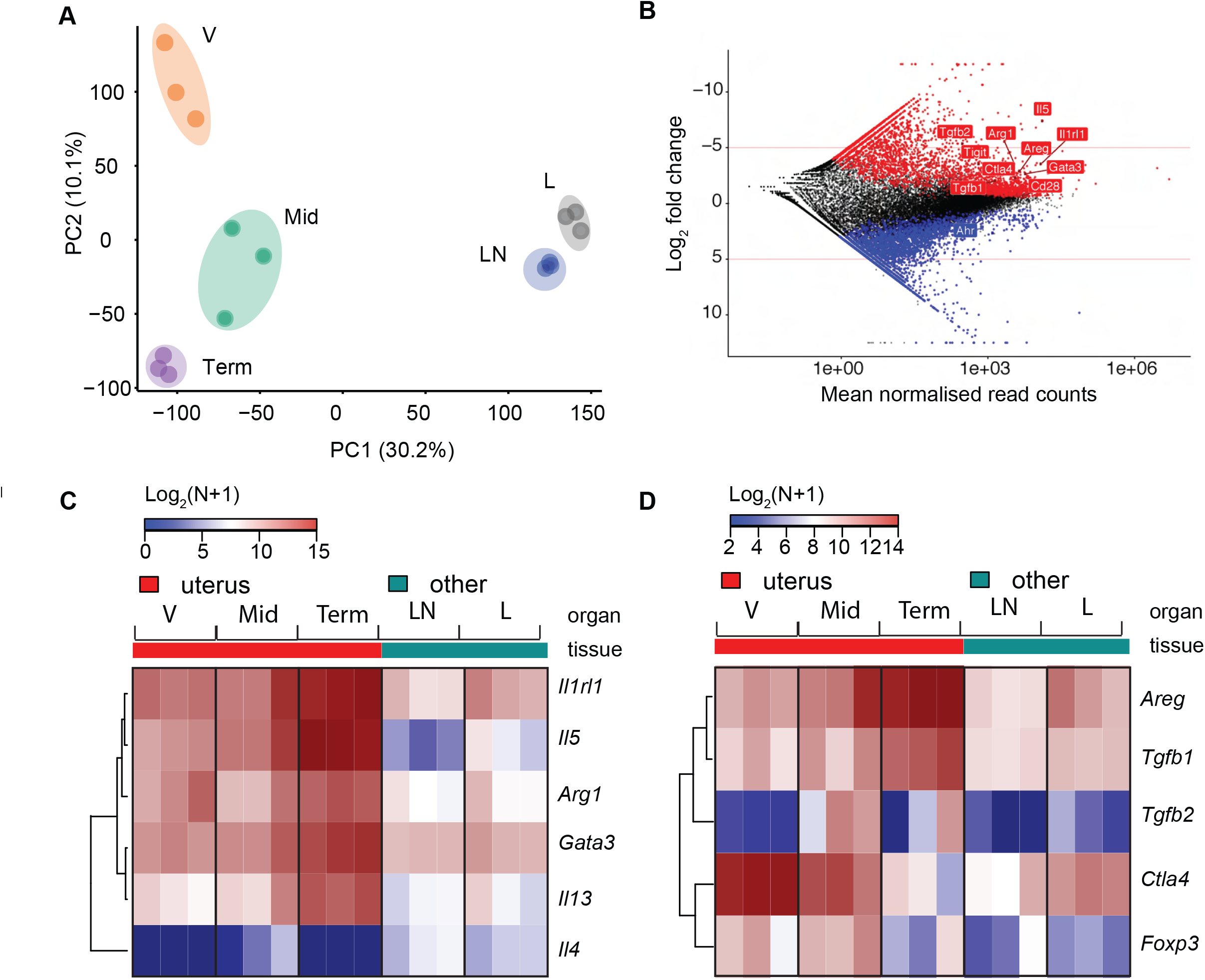
Uterine ILC2s display gene signatures of enhanced type-2 immunity. **(A)** Principal Component Analysis of RNA sequencing data from uILC2s sorted from the uterus of virgin (V) WT mice, dams from ♀WT x ♂WT crosses at mid-gestation (Mid) or term (Term), as well as from ILC2s sorted from lungs (L) and lymph node (LN) of virgin WT mice (n= 3 replicate per group); **(B)** Mean-Average plot representing differentially expressed genes between ILCs in uterus of WT virgin females or dams from ♀WT x ♂WT crosses compared to ILCs from lung (L) and lymph node of virgin WT mice (LN). Red points indicate genes higher in uterine samples and lower in L and LN samples. Genes with a significant Log2 difference less than −2-fold are the higher in uterine samples (red) or significantly greater than 2-fold are the non-uterine (red). **(C-D)** Heat maps showing normalised read count of genes identified by GO analysis as associated with (C) type-2 immunity or (D) immune regulation.

### ILC2s maintain a type-2 immune environment in the uterus

The gene signature analysis of uILC2s suggests multiple interactions with different cell types in the uterus. We therefore hypothesized that cell types interacting with ILC2s may be affected and their number altered in the absence of uILC2s. We found that, at midgestation, the uterus of ILC2KO dams had increased dendritic cells (DCs), neutrophils and macrophages, with extremely low number of eosinophils, as detected by flow cytometry (**Fig 5A, B)**. Macrophages and DCs were also visualized in uterine sections and appeared more abundant in ILC2KO dams (**Fig. S5A**). Mouse macrophages can be broadly distinguished as inflammatory M1 and alternatively activated M2 subsets, based on the expression of MHC class II molecules being higher in M1 subsets. We found that both subsets were increased in the uterus of ILC2KO dams, with a more pronounced increase in M1 macrophages (**Fig. S5B, C**). However, the numbers of uterine lymphocytes, including decidual group 1 ILC subsets (composed of tissue-resident uNK cells, conventional NK cells and uterine ILC1s) and T cells, were unaffected (**Fig. S5D-E**). This suggests that ILC2s preferentially affect the numbers of uterine myeloid cells. Notably, the anomalies in myeloid cell composition are restricted to the uterus as the numbers of DCs and macrophages in the spleen of ILC2KO dams were normal, as were the number of total splenic CD45^+^ leukocytes (**Fig 5B**). The lack of uILC2s may be expected to affect the cytokine milieu in the uterus. Strikingly, when we analysed type-2 cytokines expression in the uterus of ILC2KO dams at mid-gestation, we found that *Il4, Il5* and *Il13* transcripts were barely detectable, or much reduced, compared to WT uteri (**Fig 5C**). We also found profound skewing in gene expression profile of uterine DCs with reduced *Il4, Arg1, Chi3l3, Clec7a, Mrc1* and *Retnla* expression and modestly elevated inflammatory *Il1b* (**Fig 5D**), consistent with increased decidual *Il1b* expression in these dams (**Fig 3C**). Similar skewing was observed in uterine macrophages, with reduced *Il4, Arg1*, and *Chi3l3* expression (**Fig 5D**). In line with this finding, fewer uterine DCs and macrophages in ILC2KO dams produced Arginase-1 as detected by intracellular flow cytometry (**Fig 5E-F**), suggesting that both uterine DCs and macrophages were less polarised towards alternative activation in the absence of ILC2s. These results show that uILC2s maintain a type-2 anti-inflammatory immune environment in the pregnant uterus, including orchestrating the reprogramming of gene expression in uterine DCs and macrophages towards alternative activation to promote tissue remodeling conducive to fetal development.

**Figure 5:**
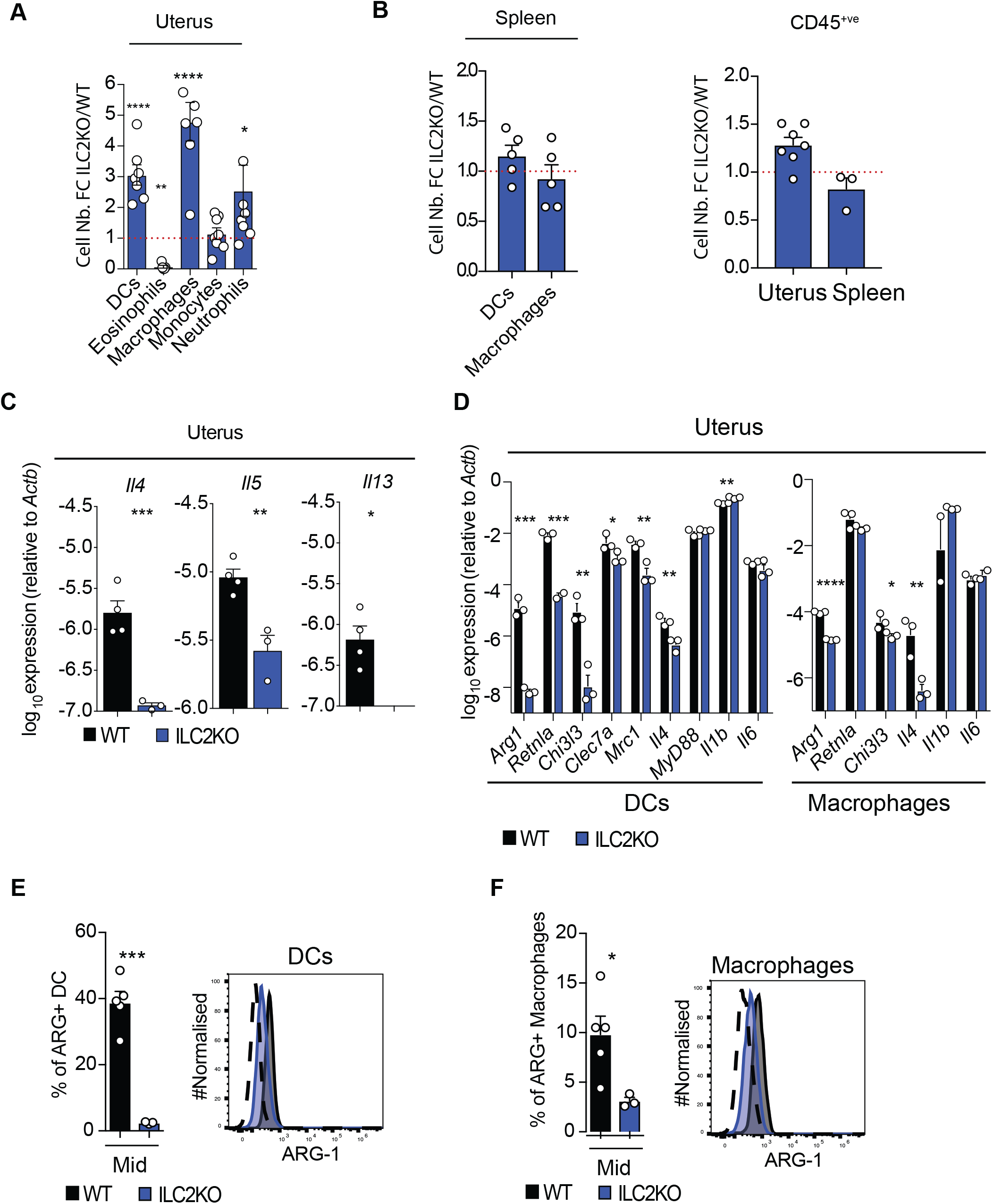
Pro-inflammatory environment and defective alternative activation of DCs and macrophages in the uterus of ILC2 KO dams. **(A)** Fold change (FC) in absolute number of indicated cell types between cells in the uterus of dams from ♀ILC2KO x ♂WT (n=7) and ♀WT x ♂WT crosses (n=10) at mid-gestation (gd9.5); **(B)** Fold change (FC) in absolute number of indicated cell types between cells in the spleen of dams from ♀ILC2KO x ♂WT (n=3) and ♀WT x ♂WT crosses (n=4) at mid-gestation (gd9.5); **(C)** Relative *mRNA* expression of type-2 cytokines in the uterus of dams from ♀WT x ♂WT crosses (n=4) and ♀ILC2KO x ♂WT crosses (n=3) at mid-gestation (gd9.5). **(D)** Relative *mRNA* expression of type-2 associated genes (*Arg1, Retnla, Chi3le, Clec7a, Mrc1* and *Il4*) and inflammatory associated genes (*Myd88, Il1b and Il6*) by purified uterine dentritic cells (DCs) and uterine macrophages of dams from ♀WT x ♂WT (n=3, black) and ♀ILC2KO x ♂WT crosses (n=3, blue) at mid-gestation (gd9.5) (n=3 pooled uteri per experiment; n=3 independent experiments). **(E, F)** Quantified (left) or representative (right) flow cytometry analysis of ARG-1 expression by purified uterine DCs (E) and macrophages (F) of dams from ♀ILC2KO x ♂WT (n=5) and ♀WT x ♂WT crosses (n=3) at mid gestation (gd9.5). Data are displayed as mean ± SEM and were analysed using Student’s *t*-test.

### ILC2-deficiency leads to enhanced endotoxin-induced abortion that correlates with expansion of IL-1β-producing uterine DC

The observed skewed immune microenvironment in the uterus of ILC2KO dams results in the observed utero-placental abnormalities and FGR (Fig. 2–3 and Fig.3 and S3). It is also possible that the effects of the dysregulated uterine micro-environment observed in ILC2KO dams may have more severe consequences during pregnancy complications such as infection. To address this hypothesis and test whether ILC2s may also protect the fetus, we administered a dose of lipopolysaccaride (LPS) that induced ~50% fetal loss in WT and *IL7ra*^*cre/wt*^ control dams, to mimic a systemic bacterial infection (**Fig. 6A** and data not shown). Notably, fetal loss was increased in ILC2KO dams, reaching over 80% (**Fig 6A, B**), suggesting a protective role of uILC2s in this system. No difference in uterine *Il6, Tnfa* and *Ifng* gene expression was detected (data not shown). However, uterine *Retnla* transcripts significantly decreased and *Il1b* increased upon LPS administration in ILC2KO dams. Interestingly, and in line with the results in Fig. 3C and Fig. 5D, both *Retnla* and *Il1b* transcripts of sham-treated ILC2KO dams were altered (**Fig 6C**). We also measured protein levels of type-2 and inflammatory cytokines in the uterus and plasma of both sham-treated and LPS-treated dams. Supporting the key role of ILC2s in maintaining a type-2 immune environment, IL-5 was significantly decreased both in the uterus and plasma of ILC2KO dams, and in both sham- and LPS-treated conditions (**Fig. S6A-B**). Uterine IL-6 was not significantly changed in either group of mice regardless of treatment. However, plasma IL-6 was increased upon LPS-treatment in ILC2KO, compared with WT dams (**Fig. S6A-B**). IL-1β was increased in both the uterus and in plasma of ILC2KO dams upon LPS challenge (**Fig. S6A-B**). Interestingly, sham-treated ILC2KO dams also showed a tendency to generate higher levels of both pro-IL-1β and mature IL-1β in the uterus compared to WT dams (**Fig. S6C**). Additionally, the number of IL-6 and TNF-α-producing macrophages in sham-treated ILC2KO dams is greater than in WT dams (**Fig S6D**). These results show increased endotoxin-induced fetal loss in pregnant mice lacking ILC2s and confirm that ILC2s sustain type-2 immunity. In the absence of ILC2s, more inflammatory cytokines are produced both locally and systemically, in response to endotoxin as well as constitutively. Thus, maternal ILC2s protect against exaggerated inflammatory responses by maintaining a balanced immune microenvironment and by preventing expansion of specific inflammatory cell types and cytokines that exacerbate pregnancy complications.

**Figure 6:**
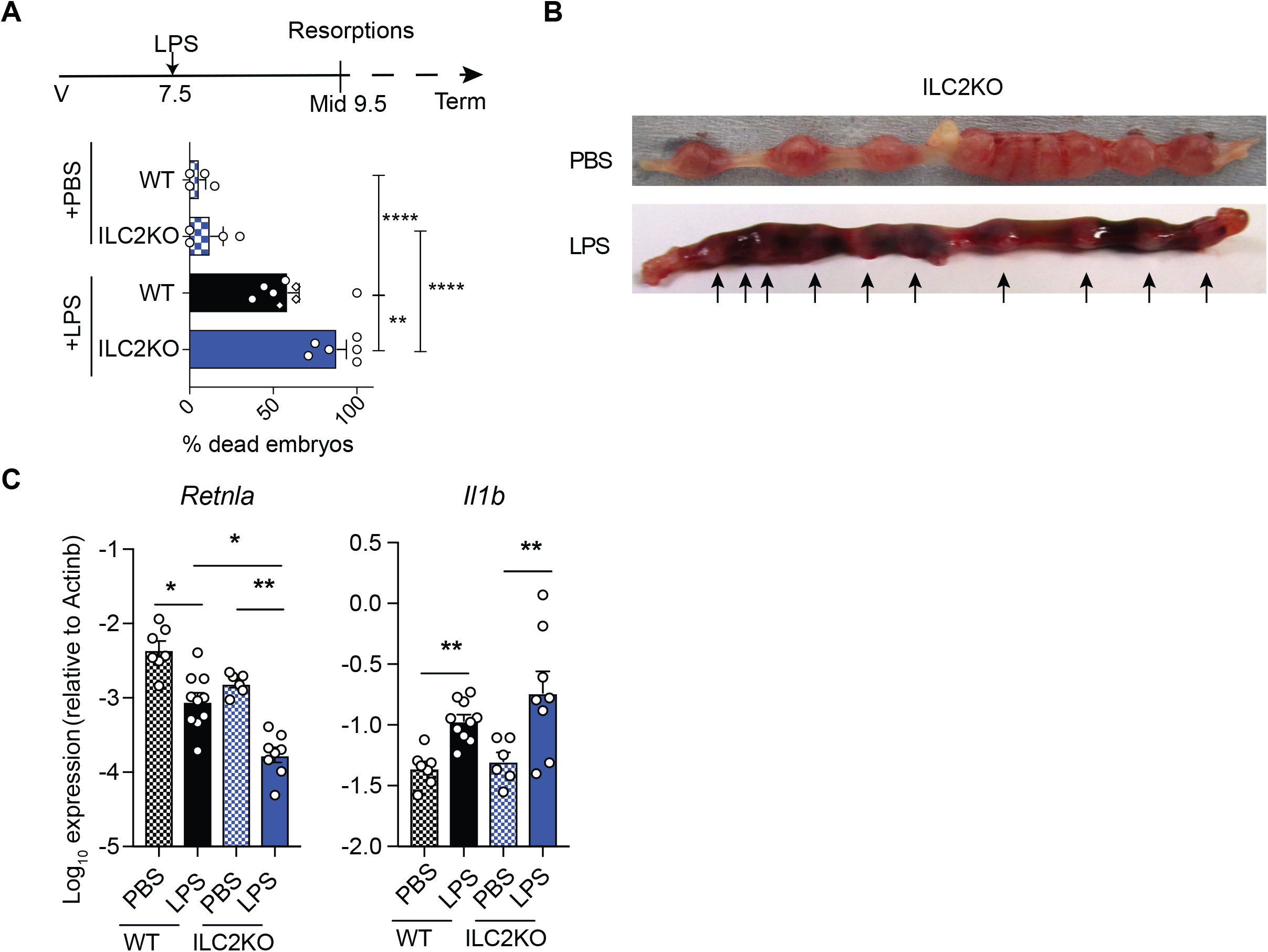
Enhanced rate of endotoxin-induced abortion in ILC2 KO dams. **(A)** Abortion rate at mid-gestation (gd9.5), 2 days post-treatment in dams from ♀WT x ♂WT crosses (sham-treated with saline, PBS, n=5; or with LPS, n=8) and ♀ILC2KO x ♂WT crosses (PBS, n=4; LPS n=6). **(B)** Representative images of the uterus of dams from a ♀ILC2KO x ♂WT cross at midgestation (gd9.5), 2 days after PBS- or LPS-treatment; **(C)** Gene expression analysis of uterine *Retnla* and *Il1b* at mid-gestation 1 day after treatment of dams from ♀WT x ♂WT crosses (PBS, n=7; LPS n=10) or ♀ILC2KO x ♂WT crosses (PBS, n=6; LPS n=8). Data in B and C are displayed as mean ± SEM. Data in B were analysed by 2-way ANOVA and data in C by Kruskal– Wallis.

### ILC2s provide a protective feedback mechanism by constraining the expansion of IL-1β-producing DCs induced by endotoxin

We hypothesised that uILC2s may counter the detrimental effects of inflammatory cytokines produced by other uterine cells, thus preventing endotoxin-induced fetal loss. Intracellular staining confirmed that a higher proportion of DCs from sham-treated ILC2KO mice expressed the inflammatory cytokine IL-1β, as compared to controls, confirming a pro-inflammatory environment prior to exposure to endotoxin (**Fig. 7A**). To dissect underlying mechanisms, we isolated leukocytes from WT mice at mid-gestation and stimulated them with LPS *in vitro*. This induced elevated numbers of ILC2s producing type-2 cytokine IL-4, IL-5 and IL-13 as determined by intracellular cytokine staining (**Fig. 7B**). It has been shown previously that ILC2s do not respond directly to LPS, as they lack TLR4. However, ILC2s can respond to IL-1β, which is released by DCs and macrophages, by increasing type-2 cytokine production (Bal et al., 2016; Ohne et al., 2016). To test if LPS-driven IL-1β-release was responsible for activating uILC2 *in vitro* we added an IL-1R antagonist to the uterine leukocyte cultures. The inhibition of IL-1 signalling completely abrogated the LPS-driven expansion of ILC2s producing type-2 cytokines (**Fig. 7B**). Consistent with the *ex-vivo* data showing LPS-driven expansion of inflammatory cells (Fig. 6 and S6), *in vitro* LPS-treated DCs from uILC2KO mice had a greater propensity to produce IL-1β (**Fig. 7C**). Notably, the addition of either IL-4 or IL-5 to LPS-treated DCs reduced the expression of IL-1β (**Fig. 7C**). Taken together, these results show that uILC2s respond to endotoxin-induced DC-derived IL-1β by increasing type-2 cytokines. In a protective feedback mechanism, these type-2 cytokines in turn suppress IL-1β-producing DCs in the uterus, and help maintain a safe environment for the fetus.

**Figure 7:**
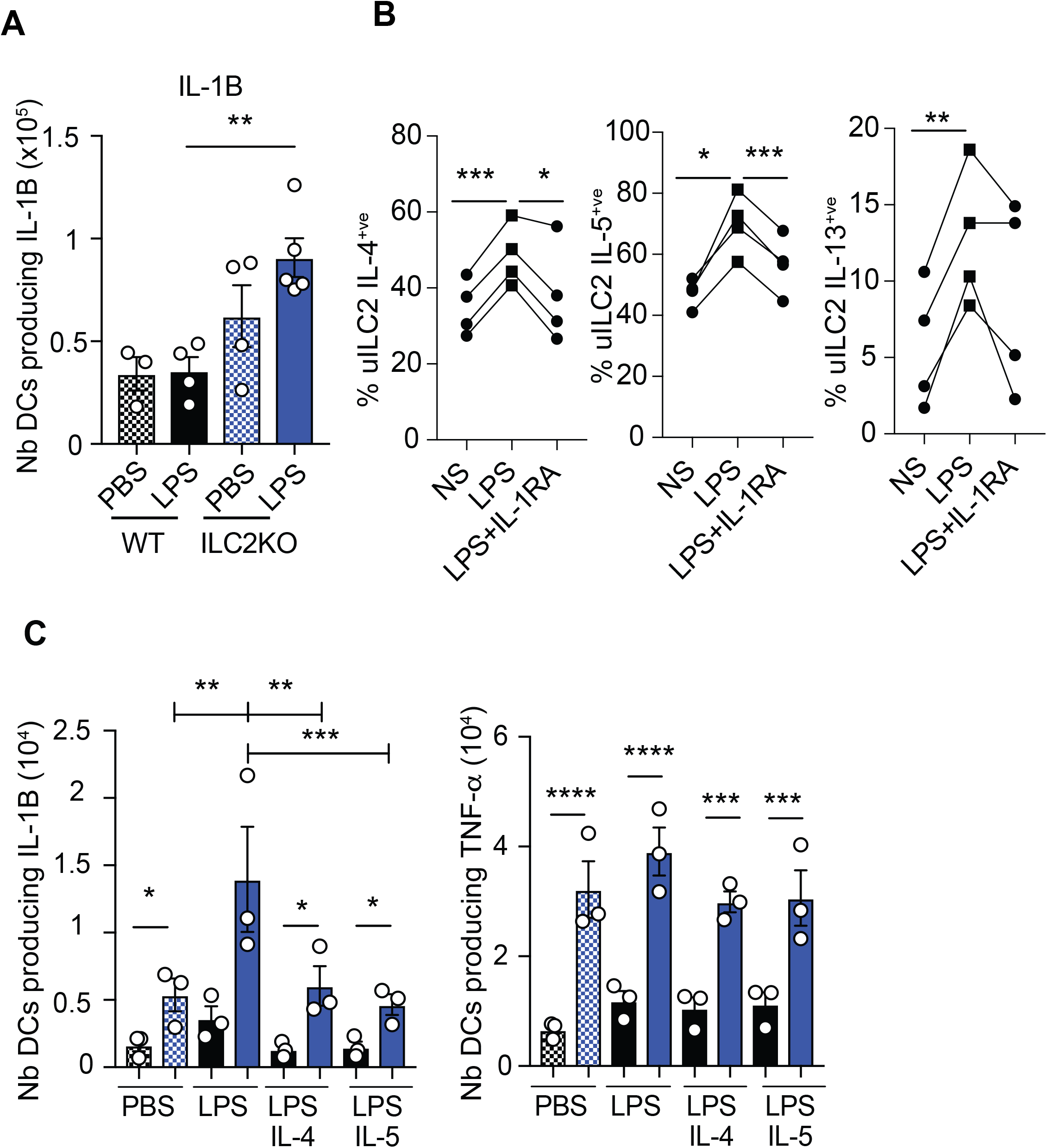
Uterine ILC2s respond to IL-1β by limiting the expansion of LPS-induced IL-1β-producing DCs. **(A)** Number of ex-vivo uterine DCs producing IL-1β at midgestation (gd9.5), 1 day after treatment of dams from ♀WT x ♂WT (PBS, n=3; LPS n=4) or ♀ILC2KO x ♂WT crosses (PBS, n=4; LPS n=5). **(B)** Leucocytes from the uterus of WT mice were stimulated in vitro by LPS for 18h, with or without IL-1R inhibitor. Percentages of uILC2s expressing the indicated type-2 cytokines are shown out of total uILC2s (n=4). **(C)** Leucocytes from the uterus of dams from ♀WT x ♂WT (n=3) or ♀ILC2KO x ♂WT crosses (n=3) were stimulated in vitro by LPS for 18h, with or without IL-4 or IL-5 and number of uterine DCs producing intracellular IL-1β (left) or TNFα (right) were counted. Data are displayed as mean ± SEM. Data in A were analysed by Kruskal–Wallis, in B with RM_one-way ANOVA and in C by RM one-way ANOVA for paired analysis and one-way ANOVA when comparing across groups.

## Discussion

Dissecting the mechanisms by which tissue resident immune cells mediate both physiological and pathological processes is one of the most exciting challenge facing modern immunologists. Our data add a new pathway to the immune regulation of pregnancy. The whole-genome transcriptome analysis comparing uterine with lung and lymph node ILC2s shows that uterine ILC2s have an enhanced type-2 immune signature. The cellular and molecular changes in the uterus of ILC2 deficient dams showed that uILC2s are crucial for the establishment and maintenance of type-2 immunity, which is required for optimal fetal growth and for safeguarding the fetus during infection (**Fig. S7**). Because the mouse model we use is not tissue-specific, we cannot exclude the effects of systemic ILC2s, although it is clear that maternal, not fetal ILC2s, contribute both to fetal growth and protection from endotoxin-induced abortion.

We have also identified an additional role for IL-1β in regulating the uILC2 response through the upregulation of type-2 cytokine production. IL-4 and IL-5 limit the expansion of IL-1β-producing DCs induced by endotoxin in a protective feedback mechanism that helps to maintain a safe environment for the fetus. Here we show that in the absence of uILC2s an acute inflammatory challenge results in elevated endotoxin-induced IL-1β production, a decreased type-2 response, and enhanced fetal loss. Since we also show that uILC2s respond to IL-1β by upregulating IL-4, IL-5 and IL-13, these results support a role for uILC2s in protecting the fetus by counteracting the effects of endotoxin-induced IL-1β and the resulting pro-inflammatory environment. While IL-4 may act directly on DCs, IL-5 may operate indirectly through tissue eosinophils (which are greatly reduced in the absence of uILC2), which in turn produce IL-4. In the adipose tissue eosinophils and ILC2s collaborate to induce the switch of macrophages from M1 to M2 (Molofsky et al., 2013). A similar mechanism may occur in the uterus, thus linking the low levels of *Il4* and eosinophils to the decrease in alternative activation in uterine DCs and macrophages observed in ILC2KO mice. It is notable that uILC2 respond to IL-1β by upregulating type-2 cytokines. This is distinct from the response reported for lung and gut ILC2s, where ILC2s exposed to IL-1β convert into IFN−γ producing ILCs, and thus contribute to the inflammatory pathology (Lim et al., 2016; Ohne et al., 2016; Silver et al., 2016). This may reflect the more profoundly type-2 environment encountered in the uterus, and the differential thresholds for mounting type-1 responses in the different tissues following pathogen exposure. Interestingly, in line with our results, human IL-4 and IL-13 downregulate LPS-induced inflammation in explanted gestation tissues (Bryant et al., 2017). Furthermore, uterine and placental deficiency in IL-4 are associated with pregnancy complications in response to inflammatory stimuli in both murine and human pregnancies (Chatterjee et al., 2014; Jiang et al., 2009; Wang et al., 2022).

The increased expression of FoxP3 and CTLA-4 in uILC2s that accompanies higher GATA-3, IL-5 and ST-2 expression, may be mediated by IL-33 (Wallrapp et al., 2017) and suggests that uILC2s may overlap with the intestinal regulatory ILCs (ILCregs) that contribute to resolution of inflammation (Wang et al., 2017). In contrast with uterine ILC2s, however, ILCregs express low levels of *Gata3, Il5* and *IL13* excluding an overlapping phenotype. It may be argued, nevertheless, that uterine ILC2s, through expressing immune regulatory molecules Foxp3 and CTLA-4, may contribute to immunological tolerance of the fetus. The traditional view on immune regulation during pregnancy is that type-2 responses are required to maintain maternal tolerance of the fetus (Offenbacher Champagne et al., 2003; Sykes et al., 2012; Wegmann et al., 1993), but KO mouse models deficient for type-2 cytokines have so far failed to show obvious phenotypes (Bonney, 2001; Fallon et al., 2002; Robertson et al., 2000; Svensson et al., 2001). However, our crosses were syngeneic and therefore not conducive to address this issue.

While we have shown the importance of the IL-33-dependent ILC2s axis in suppressing inflammatory DCs and protecting against endotoxin-induced abortion, precisely how ILC2s regulate placental function and fetal growth remains to be determined. The failure to achieve the genetically determined growth potential in fetuses may be due to inadequate maternal or placental responses to fetal demands. Our data clearly show that fetal growth restriction is linked to the maternal genotype and that maternal ILC2s influence the utero-placental unit. Other maternal ILC subsets, notably uNK cells, have clearly been shown to regulate the utero-placental unit, with consequences on fetal and placental growth (Gaynor & Colucci, 2017). The feto-placental phenotype of dams with hypofunctional or reduced uNK cells partially overlaps with that of ILC2-deficient dams, including incomplete uterine decidual adaptations, placental insufficiency and fetal growth restriction (Ashkar et al., 2000; Boulenouar et al., 2016; Kieckbusch et al., 2014). In addition, ILC2-deficient dams show spontaneously increased *Il1b* and decreased *Il4, Il5* and *Il13* transcripts in the uterus, together with cell-intrinsic gene signatures consistent with polarization towards inflammatory responses in resident uterine DCs and macrophages, alongside dysregulated immune cells consistent with a pro-inflammatory environment. Accordingly, the endometrium of women with recurrent pregnancy loss shows features of a pro-inflammatory microenvironment with reduced tolerogenic DC (Guo et al., 2021; Liu et al., 2018).

IL-1β is a highly pleiotropic cytokine that may play multiple roles in pregnancy and *IL1B* polymorphism is linked to miscarriage (Wang et al., 2022; Zhang et al., 2017). The poor decidual arterial remodelling and placental transport capacity observed in ILC2 deficient dams can be caused by elevated IL-1β levels and is expected to reduce blood supply of resources, subsequently leading to reduced fetal growth in later pregnancy. Although IL-1β does not cross the placenta, it can directly affect the invading fetal trophoblast. Maternal or placental IL-1β is known to reduce insulin signalling in human trophoblast (Aye et al., 2013). The nature of fetal signals that regulate placental allocation of maternal resources is largely unknown (Constância et al., 2002; Higgins et al., 2016; Jansson, 2016). Fetal genes do regulate fetal growth (Warrington et al.,2019), however maternal genotype also influences placental functions and fetal growth (Kieckbusch et al., 2015; Sferruzzi-Perri et al., 2016). It is conceivable that fetal growth restriction in ILC2KO dams might have been even more marked in the absence of placental adaptations later in pregnancy, such as increased expression of transporter genes and greater maternal blood spaces. Similar compensatory strategies are seen in other adverse gestational environments, like undernutrition, hypoxia and obesity(Constância et al., 2002; Higgins et al., 2016). This may be the first study to describe placental adaptation to an immune-mediated stress.

Growth restricted fetuses are associated with asymmetric growth because a circulatory adaptation preferentially shunts blood to the head, preserving its growth. ILC2KO dams produced symmetric fetal growth restriction, with no apparent brain sparing. The lack of brain sparing may be due to the low degree of growth restriction, which may be relatively well tolerated and therefore does not trigger the circulatory adaptation. Future work will be required to dissect the mechanisms and assess the extent of defective fetal growth in ILC2 KO dams.

Despite these anomalies in ILC2KO dams, they have litters of normal size with low neonatal mortality. Therefore, the immune dysregulation found in ILC2KO dams seems to be relatively well tolerated under pathogen-free conditions and in steady state. By contrast, increased neonatal mortality has been reported in pups from ST-2 KO dams (Bartemes et al., 2017), suggesting that maternal IL-33 signaling may affects additional pathways that also impact on neonatal survival.

In conclusion, IL-33 dependent maternal ILC2s emerge as key cells in maintaining a healthy utero-placental environment conducive to both fetal growth and protection against endotoxin-induced abortion (**Fig. S7**). Our data establish that maternal ILC2s are critical for the achievement of genetically determined fetal growth potential and for safeguarding the fetus by preventing abortion induced by systemic endotoxin insult.

## Acknowledgements

We thank the members of the Colucci and McKenzie labs and Tim Halim for suggestions and discussions. This work was funded by the Centre for Trophoblast Research, the Wellcome Trust (Grant 200841/Z/16/Z to FC and 100963/Z/13/Z to ANJM), the Medical Research Council (U105178805 to ANJM) and the Cambridge NIHR BRC Cell Phenotyping Hub to FC. We thank Jan Brosens for providing human endometrial biopsies, Hans-Reimer Rodewald for providing IL7RαCre mice, Esther Perez, Natalia Savinykh and Christian Bowman for cell sorting, the staff at the CBS and ARES animal facilities, particularly Nicola Jeyes and Shona Butler, Helen Jolin and Alexandros Englezakis for help in ARES animal facility, and the Metabolic Research Laboratories (MRL) CBAL, GTU and histology facilities. EB was supported by a CTR PhD fellowship and ANSP by a Royal Society Dorothy Hodgkin Fellowship. The authors declare no competing financial interests. All authors approve the final version of the manuscript.

## Figure Legends

**Figure S1: Phenotype and frequencies of murine and human uterine ILC2s before and during pregnancy**.

**(A)** Diagram showing tissue localisation of uterine (u) macrophages (blue), dendritic cells (DCs, orange), NK cells (red) and ILC2s (green) in virgin WT mice, and in dams from ♀WT x ♂WT crosses at mid-gestation (gestation day 9.5 = gd9.5) and at term (gd18.5). The mesometrial lymphoid aggregate of pregnancy (MLAp), that forms in dams between the two uterine smooth muscle layers is highlighted. Representative flow cytomery data show CD45^+ve^ NK1.1^-ve^ NKp46^-ve^ CD11b^-ve^ CD3^-ve^ CD19^-ve^ CD90^+ve^ CD127^+ve^ uILC2s at each time point during pregnancy. Absolute numbers of uILC2s are indicated above each plot. Data are displayed as mean ± SEM (n=8 pools of 10 uteri from WT virgin mice, 10 uteri from WT dams at mid-gestation and 4 uteri from WT dams at term). (**B**) Flow cytometry analysis of *Il33* expression in the mouse uterus, which was restricted to the CD45^-ve^ fraction in heterozygous *Il33*^*cit/wt*^ dams <u>at mid-gestation</u> and **(C)** is present in the myometrium (M) but not detectable in the decidua (Dec) (n = 3 biological replicates). **(D)** IL-33 protein from uterine tissue homogenates of dams from ♀WT x ♂WT crosses at midgestation (gd9.5) as detected by ELISA (n=3 biological replicates). **(E)** Gating strategy defining human uILC2s as detected by flow cytometry of cell suspensions from endometrium (non-pregnant) and decidua of pregnancies voluntarily terminated between gestation week 7 and 12. Histograms show human uILC2s (green), Lin^-ve^, CD56^+ve^, CD9^+ve^ uNK cells (red), Lin^−^ CD127^+^, ROR-γt^+^, CD56^low^ uILC3s (blue) and CD117^+^ CRTH2^+^ uILC2s (pink). **(F)** Percentage of human uterine ILC2s from endometrium obtained from biopsies 6-10 days after the pre-ovulatory luteinizing hormone surge and decidua obtained from elective pregnancy termination at 7-12 weeks of gestation (n=10 biological replicates). Data in C, D and F are displayed as mean ± SEM.

**Figure S2: Decreased placental efficiency but normal litter size from ILC2 KO dams**.

**(A)** Number of new born (left) and neonatal mortality rate before weaning (right) from ♀WT x ♂WT crosses (n=107 pregnancies from 20 dams) and ♀ILC2KO x ♂WT crosses (N=60 pregnancies from 17 dams). **(B)** Ratios between fetal and placental weights from ♀WT x ♂WT crosses (n=13 litters); ♀ILC2KO x ♂WT crosses (n=8 litters), ♀*Il33*^*cit/cit*^ x ♂WT crosses (n=4 litters), ♀WT x ♂ILC2KO crosses (n=7 litters). Data in A and B are displayed as mean ± SEM. Data in A were analysed by Student’s *t*-test. Data in B were analysed by mix-model approach (accounting for inter-litter variability but not sex). **(C)** Ratios between brain volume and fetal weight. Data from Figure 2 were used to calculate the ratio between fetal brain volume and fetal body weight in gd18.5 fetuses from ♀ILC2KO x ♂WT crosses (n=9 from 3 dams) and ♀WT x ♂ILC2KO crosses (n=12 from 4 dams).

**Figure S3: Increased maternal blood spaces in the placentas of ILC2 KO dams**.

Quantification of maternal blood spaces in sections of placentas at term (gd18.5) stained for lectin, cytokeratin and eosin, as shown Figure 3E (n=4 placentas/group). Data are displayed as means ± SEM and were analysed by Student’s *t*-test.

**Figure S4: Enhanced expression of IL-5, ST-2, CTLA-4, and Foxp3 in uterine ILC2s**.

**(A, B)** Representative (left) and quantified (right) flow cytometric analysis of (A) CTLA-4 and (B) FOXP-3 expression by lung (grey) and uterine (green) ILC2s from dams of ♀WT x ♂WT crosses at mid-gestation. **(C)** Flow cytometry analysis of *in vitro* cultured lung (grey) and uterine (green) ILC2s from dams of ♀WT x ♂WT crosses at mid-gestation treated with PBS, IL-33 and IL-25 or PMA-ionomycin (P/I). uILC2s expression of IL-5, IL-13, ST-2 and AREG expression are shown as fold change relative to that of lung ILC2s. (n=5 uteri per experiment, 3 independent experiment per time point). Data are displayed as mean ± SEM and were analysed by Student’s *t*-test. **(D)** Heat maps showing normalised read count of genes in ILC2s from the uterus of dams from ♀WT x ♂WT crosses at mid-gestation, compared to ILC2s from “other” (lung and lymph node) tissues.

**Figure S5: Macrophages, DCs and lymphocytes in the uterus of ILC2 KO dams**.

**(A, B)** Immunohistochemical analysis of (A) uterine macrophages identified as F4/80 positive (brown staining) and (B) DCs identified as MHC-II positive (red staining) and CD11c positive (brown staining) in dams from ♀WT x ♂WT or ♀ILC2KO x ♂WT crosses at midgestation (gd9.5). 14X magnification (top) and 4X magnification (bottom). **(C)** Fold change (FC) in the number of MHC-II high (type-1 polarised) and MHC-II low (type-2 polarised) macrophages in the uterus of dams from ♀WT x ♂WT (n=10) or ♀ILC2KO x ♂WT crosses (n=7) at midgestation (gd9.5), as determined by flow cytometry. **(D)** Fold change in the number of group 1 ILCs, including tissue resident NK cells (trNK), conventional NK cells (cNK) and uterine ILC1s (uILC1) and T cells (left) and relative composition of group 1 ILCs in the uterus of dams from ♀WT x ♂WT (n=3) or ♀ILC2KO x ♂WT crosses (n=3) at midgestation (gd9.5), as determined by flow cytometry. Data are displayed as mean ± SEM and were analysed by unpaired Student’s *t*-tests.

**Figure S6: Increased inflammatory response in ILC2 KO dams**.

Cytokines detected by Multiplex ELISA in **(A)** uterine tissue homogenates or **(B)** plasma from dams at mid-gestation, 1 day after systemic administration of PBS or LPS; ♀WT x ♂WT (PBS, n=4; LPS n=6); ♀ILC2KO x ♂WT (PBS, n=3; LPS n=3) (**C)** Identification and quantification by western blot of both active and pro-IL-1β protein from uterine homogenates collected at midgestation (gd9.5) 1 day after treatment of dams from ♀WT x ♂WT (PBS, n=3; LPS n=5) and ♀ILC2KO x ♂WT crosses (PBS, n=2; LPS n=4). Gel lanes cut from the original western, which is shown separately in Supplementary Figure S8 (lane numbers displayed are 1, 5, 9 and 14). **(D)** Number of ex-vivo uterine DCs (left) and macrophages (right) at midgestation (gd9.5) producing IL-6 or TNF-α, 1 day after treatment of dams from ♀WT x ♂WT (PBS, n=3; LPS n=4) and ♀ILC2KO x ♂WT crosses (PBS, n=4; LPS n=5). Data in A, B and D are displayed as mean ± SEM and were analysed by one-way ANOVA.

**Figure S7: Summary diagram of the proposed mechanism**

Diagram summarising the molecular mechanism of the roles ILC2s play during physiological pregnancy and bacterial infection. In normal pregnancy, uILC2s sustain the production of type-2 cytokines to favour type-2 polarisation of myeloid cells. This controlled immune environment promotes normal decidual vascularization, placental function and fetal growth. In murine models lacking ILC2s, these processes are compromised leading to FGR, including reduced brain volume. During systemic endotoxin challenge, LPS activates DCs and macrophages in the uterus, which produce inflammatory cytokines (including IL-1β), which in turn stimulate uterine ILC2s to produce more type-2 immune cytokines, thus contrasting IL-1β-producing DCs and protecting against excessive inflammation. In the absence of maternal ILC2s in pregnant KO females, the excess of uterine myeloid cells and their inflammatory cytokine production cannot be counterbalanced by prompt production of type-2 cytokines, resulting in enhanced rate of abortion.

## Materials and Methods (2856-also the ref counts)

### Mice

C57BL/6 (B6) WT mice were purchased from Charles River. *Il33*^*cit/wt*^, *Il33*^*cit/cit*^(Hardman et al., 2013) *Rora*^*flox/flox*^ *Il7ra*^*cre/wt*^ (Oliphant et al., 2014) and *Il7ra*^*cre/wt*^ (Schlenner et al., 2010) were all on B6 background and maintained at the University of Cambridge Central Biomedical Service or Medical Research Council (MRC) ARES animal facilities, under pathogen-free conditions according to UK Home Office guidelines. Mice were used at 8-12 weeks of age. In both (♀ILC2KO x ♂WT) and (♀WT x ♂ILC2KO) crosses, fetuses and placentas are genetically equivalent for *Rora* and express either one or no copy of the *Il7ra*^*cre*^ transgene. Virgin mice were at random stages of estrous cycle.

### Human tissues

Decidual biopsies were obtained from donors undergoing elective termination between 7 and 12 weeks of pregnancy. This study was approved by the Cambridge Research Ethics Committee approved (04/Q0108/23 and 08/H0305/40) and the NHS National Research Ethics – Hammersmith and Queen Charlotte’s & Chelsea Research Ethics Committee (1997/5065). Endometrial biopsies were timed between 6 and 10 days after the pre-ovulatory luteinizing hormone surge and were obtained in ovulatory cycles. Subjects were not on hormonal treatments for at least 3 months prior to the procedure. Subjects were recruited from the Implantation Clinic at University Hospitals Coventry and Warwickshire National Health Service Trust. Written informed consent was obtained from all participants in accordance with the guidelines in The Declaration of Helsinki 2000. Human decidua and endometrium tissues were mechanically processed and leukocytes were enriched by layering on Lymphoprep (Axis-Shield).

### Murine tissue preparation

Lymph nodes were passed through a 70μm cell strainer using PBS 2% FCS. For lung digestion to obtain samples for RNAseq analysis only, lungs were mechanically dissociated in RPMI-1640, and digested with collagenase I (Life Technologies), DNase I (Roche) at 37°C whilst shaking. In all other experiments mouse uteri, spleen, and lungs, were processed using two different enzymatic protocols to obtain either leukocytes or myeloid cells and stromal cells. For leukocytes, tissue were subjected to mechanical dissociation followed by incubation with HBSS (Ca^++^ and Mg^++^ free Life Technologies), 10% FCS (Life Technologies), 5 mM EDTA (Sigma), 15 mM HEPES solution (Life Technologies) on a rotator at 37°C. Samples were then digested in RPMI 2% FCS, 30 μg/ml DNase I (Roche) and Liberase DH 0.1 WU/ml (Roche). Next, tissues were passed through a 100μm cell strainer and leukocytes were enriched using a 80%/40% Percoll (GE Healthcare Life Sciences) gradient. Myeloid cells and stromal cells were obtained by incubating minced tissues with HBSS containing Ca^++^ and Mg^++^ (Life Technologies), 30 μg/ml DNase I (Roche) and 0.26 WU/ml Liberase TM (Roche) on a rotator at 37°C then incubated with 5mM EDTA (Sigma) in PBS. Cell suspensions were filtered using a 100μm cell strainer.

### *In vitro* cell activation

Uterine tissues were harvested at gd9.5 and cell suspensions from each mouse were equally divided per treatment, to allow the direct comparison of different treatments on the same biological replicate. For LPS stimulation, cells were incubated overnight with 5μg/ml LPS (LPS *E coli* 055:B5 Sigma), with brefeldin A and monensin cocktail (Biolegend) for the last 4h of culture, in the presence or absence of recombinant IL-1Rα (R&D 100ng/ml). Alternativelly to IL-1Rα? recombinant murine IL-4 (10ng/ml Peprotech) or IL-5 (10ng/ml Peprotech) or PBS were added to the culture to assess the importance of type 2 cytokines during LPS response. For intracellular cytokine detection cells were activated using PMA/ionomycin activation cocktail (Biolegend) for 4 h or with 20 ng/ml rmIL-25 and 20 ng/ml rmIL-33 for 4 h in the presence of brefeldin A and monensin cocktail (Biolegend). For IL-4 detection cells were incubated in the presence of brefeldin A and monensin cocktail (Biolegend) only for 4 h.

### Flow cytometry and cell purification

Human leukocytes purified from decidua, endometrium or myometrium were stained with antibodies specific for the following human antigens: CD45 (HI30), CD3 (UCHT1), CD19 (HIB19), NKp44 (P44-8), CD14 (M5E2), CD9 (M-L13), CRTH2 (BM16), CD56 (HCD56), CD117 (104D2), CD127 (A019D5), CD161 (HP-3G10), GATA3 (TWAJ and 16E10A23), T-BET (O4-46) and RORγt (Q21-559). Murine cells were stained with antibodies specific for the following mouse antigens: CD45 (30-F11), CD3 (17A2), CD4 (GK1.5), CD8a (53-6.7), CD11b (M1/70), CD11c (HL3), CD19 (6D5), F4/80 (BM8), ICOS (C398.4A), NK1.1 (PK136), NKp46, CD90.2 (53-2.1), CD127 (A7R34), ST-2 (RMST2-33), CD49a (Ha31/8), MHC II I-A/I-E (M5/114.15.2) EOMES (Dan11mag), TER-119 (TER-119), IL-1β (166931 R&D), IL-5 (TRFK5), IL-6 (MP5-20F3) IL-13 (eBio13A), TNFα (MP6-XT22) and Areg (AF989), GATA3 (TWAJ and 16E10A23), T-BET (O4-46) and RORγt (Q21-559). CD80 (16-10A1), SiglecF (E50-2440), Ly6C (AL-21), Ly6G (1A8), GR-1 (RB6-8C5), ARG-1 (Sheep IgG Polyclonal), CD31a (390), Podoplanin (8.1.1), SCA-1 (D7), CD34 (RAM34), PDGFRA (APA5), RELMα (Rabbit polyclonal Peprotech). AREG and RELMa antibodies were conjugated in house by using lightning-link conjugation kit (Innova Biosciences). All antibodies were purchased from Biolegend, eBioscience, BD Biosciences, Peprotech or R&D Systems. DAPI, Fixable viability dye eFluor 780, eFluor 450 and eFluor 506 (eBioscience) were used to exclude dead cells. Cytokines and transcription factors were stained using the FoxP3 staining buffer set (eBioscience) according to the manufacturer’s instructions. Samples for immune phenotyping were acquired on LSR Fortessa (BD Biosciences) using FACS DiVa and analyzed using FlowJo (Tree Star). All hematopoietic cells were gated as live single CD45^+^ cells. Cell populations were identified as follows; uDCs; MHC II^high^ F4/80^−/low^ Ly6g^−ve^ Ly6c^− ve^ CD11c^+^; uMac as MHC II^+^ F4/80^+^ Ly6g^−ve^ Ly6c^−ve^ CD11b^+^; uNK cell as F4/80^−ve^ Ly6g^−ve^ Ly6c^−ve^ CD3^−ve^, NK1.1^+^ and uT cells as F4/80^−ve^ Ly6g^−ve^ Ly6c^−ve^ NK1.1^−ve^ CD3^+^CD4^+^. The following populations were fluorescence-activated cell sorted (FACS) for RNAseq analysis; ILC2s from pooled mesenteric, para-aortic and inguinal lymph nodes or the lung were flow-sorted as lineage^− ve^ CD127^+^ ICOS^+^ and uILC2s as live CD45^+^ CD11b^−ve^ NK1.1^−ve^ CD3^−ve^ CD19^−ve^ GR-1^−ve^ Ter119^−ve^ CD11c^−ve^ CD8a^−ve^ ST2^+^ CD90^+^ from virgin mice, as well as uILC2 from dams at gd9.5 and gd18.5. Cell were sorted using a Sony iCyt Synergy or BD FACSAria III to >95% purity.

### Gene expression analysis

RNA extractions were performed using TRI Reagent (Sigma) purified using RNAeasy mini kit (Qiagen). gDNA digestion was performed using Turbo DNAse (Ambion). RNA was re-purified and concentrated using the RNAeasy micro kit (Qiagen). RNA from homogenised uterine myometrium were isolated with TRI Reagent (Sigma) in Lysis matrix D (MP Bio). Alternatively, RNA from LPS treated samples were performed on whole tissues using TRI Reagent (Sigma) in Lysin matrix D (MP Bio) followed by chloroform extraction (Sigma) and purification using RNAeasy mini kit (Qiagen). RNA integrity was assessed by Picochip (Agilent) using a Bioanalyzer 2100 (Agilent). For RNA sequencing library preparation was performed using NuGEN Ovation RNA-seq system V2 and NuGEN Ovation Ultralow DR Multiplex System kit and assessed by high sensitivity DNA chip usgin a 2100 Bioanalyzer (Agilent). RNA sequencing was performed on a Illumina HiSeq2500. RNAseq data analysis was then performed as follows. Raw FASTQ files were adapter, and quality trimmed using trim_galore(Babraham Bioinformatics, n.d.-b) (v0.4.1). Quality control was performed with FastQC (v0.11.5)(Babraham Bioinformatics, n.d.-a). Data were aligned to GRCh38 mouse genome (Ensembl Release 86) with TopHat2 (v2.1.1, using bowtie2 v2.2.9)(Kim et al., 2013) with a double map strategy. Alignments and QC were processed using custom ClusterFlow (v0.5dev)(Zissler et al., 2015) pipelines and assessed using MultiQC (0.9.dev0)(Ewels et al., 2016). Gene quantification was determined with HTSeq-Counts (v0.6.1p1)(Anders et al., 2015). Additional quality control was performed with feature counts (v 1.5.0-p2)(Liao et al., 2014), Qualimap (v2.2)(García-Alcalde et al., 2012) and Preseq (v2.0.0)(Daley & Smith, 2013). Differential gene expression analysis was performed and read counts were normalised on the estimated size factors using with DESeq2 package (v1.16.1, R v3.4.0)(Love et al., 2014). Custom scripts are available: https://github.com/CTR-BFX/2017-Balmas-Colucci. Gene Ontology analysis was performed using DAVID 6.8 by uploading gene lists and their directional fold changes. Gene Ontology Lists were then filtered and only GO lists with at least 4 genes and with significant p-value were considered and ranked per significance. EMBL-EBI Accession number reference: E-MTAB-5803 and E-MTAB-5806. For real-time PCR analysis, RNA was and reverse transcribed using a high capacity RNA-to-cDNA kit (Applied Biosystem) and qPCR was performed using Power SYBR green (Thermo Fisher) with gene specific primers (see **Table S2**) using a Quant Studio 6-flex real time PCR (Thermo Fisher). 2^ΔCt^ were calculated per every gene as relative to *Actb*. Primers previously shown to exhibit efficiencies > 95% were selected.

### Cytokine detection

Uterine tissues from virgin mice or dams (myometrium and decidua at mid-gestation and whole uterus at term) were homogenised using a tissue homogeniser (Stuart) with 1X cOmplete EDTA free protease inhibitor tablets (Roche) and total protein was quantified using a Pierce BSA assay (Thermo Fisher). IL-33 was examined using a Mouse IL-33 Ready-SET Go! (eBioscience) ELISA and data were normalized to total protein. IL-1β was detected by westernbot analysis. 10μg per sample were loaded and separated on Any kD Mini-PRO-TEAN TGX precast polyacrylamide gels from Bio-Rad (Hercules, CA). Proteins were transferred onto PVDF membranes (Thermo Scientific) and blocked with 5% BSA for 1h at room temperature. After washing in Tris-buffered saline containing 0.1% Tween (TBS-T), membranes were incubated with IL-1β antibody (#12242 CST, Danvers MA) overnight at 4°C in TBS-T containing 2.5% BSA. The membranes were washed and incubated with peroxidase-conjugated anti-rabbit antibody (1:2000; #7074 CST) diluted in 2.5% BSA and visualised by enhanced chemiluminescence (ECL) using ECL Western Blotting Substrate on the ChemiDoc XRS system (BioRad). Densitometry analysis was performed using Image J software (imagej.nih.gov). Target protein levels were normalised to total protein levels as determined by Amido Black (Sigma-Aldrich) staining.

### Immunohistochemistry

Uterine tissues were dissected at gd9.5 and fixed in 10% formalin for 5h at RT. Tissues were paraffin-embedded and cut into serial sections at 7μm. To assess arterial remodeling sections were stained in 50% Hematoxylin Gill (Sigma) and Eosin (Sigma) and evaluated as described in Kieckbusch et al, Jove, 2015(Kieckbusch, Gaynor, et al., 2015).To assess macrophage infiltration antigen retrieval with either proteinase K (Sigma) or sodium citrate buffer was perfomed followed by avidin-biotin blocking (Vector Labs), endogenous peroxidase activity was blocked in 3% H_2_O_2_ in ddH_2_O to avoid non-specific staining. Macrophages were stained with Rat IgG2b anti-mouse anti F4/80 (MCA497A488T BioRad) with Immpress Kit (Vector labs). To detect citrine mouse anti-mouse anti YFP (Abcam) and MOM kit (Vector Labs) combined with Vectastain Elite ABC-Alkaline phosphatase kit (Vector Labs) were used. To stain for DCs sections in cryo-embedded uterine tissue from pregnant females at gd9.5 were cut at 7μm and fixed in 4% formalin. Rat IgG2b anti-mouse anti MHC I-a/I-e (Biolegend) was used in combination with impress-AP (Vector Labs) and Hamster anti mouse CD11c (Biolegend) was used in combination with biotinylated secondary anti-hamster IgG (Vector Labs) and Vectastain Elite ABC-HRP kit (Vector Labs). We used VectorRed (Vector Labs) and SigmaFast DAB (Sigma) as Alkaline Phosphate and Peroxidase substrate. To assess placental morphology placentas were dissected at gd18.5, processed as described above and stained with haematoxylin and eosin. Absolute and percentage volumes of the placental labyrinthine and junctional zones (Lz and Jz respectively) as described previously(Coan et al., 2004; Sferruzzi-Perri et al., 2016) was determined by point counting using the Computer Assisted Stereological Toolbox (CAST v2.0; Olympus). To assess Lz composition and component volumes, placental sections were double-labelled with rabbit antibodies against cytokeratin (Thermo-Fisher Scientific 180059) and lectin (Vector Labs B-1205) to identify trophoblast and fetal capillaries respectively, and counterstained with haematoxylin and eosin. For lectin visualisation antigen retrieval with 10mM sodium citrate buffer was performed, endogenous peroxidases were quenched and samples were blocked using 1% bovine serum albumin (BSA) in PBS and incubated overnight with lectin antibody (1:50 dilution in PBS) at 4°C. Streptavidin-horse radish peroxidase complex (Strep-HRP Rockland S000-03, 1:250 in PBS) was then applied and Lectin was detected by staining with diaminobenzidine (Sigma) in saturated ammonium nickel (II) sulfate solution. Cytokeratin staining was perfomed similarly with the following modifications. Cytokeratin antibody (1:100 in 10% goat serum and 1%BSA/PBS), with an additional 1h incubation step with goat anti-rabbit secondary antibody at room temperature was performed before Strep-HRP was added, nickel in the chromogen was omitted. Sections were dehydrated in increasing ethanol concentration and mounted with DPX. Omission of the primary antibodies was used as the negative control. Point counting was used to evaluate the fetal capillary (FC), trophoblast and maternal blood space (MBS) volume densities within the Lz, which were multiplied by the estimated Lz volume to acquire estimated volumes of individual components. Twenty random fields of view (320 points) were scored per placenta. MBS and FC surface densities were determined by counting the number of intersecting points along cycloid arcs in random fields of view. Multiplying these values by the estimated Lz volume provided the absolute surface areas, which averaged gives the total surface area available for exchange. Harmonic barrier thickness was estimated by measuring the shortest distance between FC and the closest MBS, calculating the mean of the reciprocal of those measures and then correcting for the plane of sectioning by a factor of 0.85. A total of 200 measurements were made for each Lz. Theoretical diffusion capacity was calculated by dividing the total surface area for exchange by the barrier thickness and multiplying by Krogh’s oxygen diffusion constant. Specific diffusion capacity was estimated by dividing theoretical diffusion capacity by the fetal mass. FC length densities per section were determined using counting frames and then converted to FC length and diameter. For citrine (*Il33*) detection in gd9.5 uteri, tissues were fixed in phosphate-buffered 1% formaldehyde (methanol-free, Thermo Fischer Scientific), transferred to 30% sucrose, embedded in 15% sucrose + 7.5% porcine skin gelatin (Sigma) in PBS and flash-frozen in isopentane −80°C. 10 μm sections were mounted on onto Superfrost Plus slides (Thermo Scientific) blocked using 1% donkey serum (Jackson Immunoresearch) in 0.05% Triton-X in PBS and citrine was detected using polyclonal rabbit anti-GFP antibody (Life Technologies) (5 μg/ml) and alexa Fluor 488-labeled polyclonal donkey anti-rabbit antibody (Life Technologies) (4 μg/ml). Nuclei were stained using DAPI (300 nM). Prolong Gold (Invitrogen) was used to mount coverslips. Sections were imaged using a Nikon HCA.

### Evaluation of fetal survival

Breeding records from *Rora*^*flox/flox*^ *Il7ra*^*cre/wt*^ were analysed to assess fetal loss in ILC2KO mice. We compared the number of newborn and fetal survival in *Rora*^*flox/flox*^ *Il7ra*^*cre/wt*^ dams mated with ILC2 competent sires (♀ILC2KO x ♂WT cross) and in ILC2 competent dams mated with ILC2 competent sires (♀WT x ♂WT cross). Therefore, the pups are ILC2 competent in both groups. Breeding records from *Rora*^*flox/flox*^ *Il7ra*^*cre/wt*^ were analysed with GraphPad Prism by unpaired Student *t*-test to assess foetal loss in ILC2KO mouse models.

### Fetal weight, placental weight and foetal survival

Fetuses and placentas from gd18.5 were weighed and fetuses were harvested into 4% PFA and hydrogel embedded for CT scan analysis. Pups from *Il33*^*cit/cit*^, *Il33*^*cit/+*^ or WT mothers mated with WT males, were weighed at day 7 and day 14 after birth. Statistics on newborn weights, fetal weights, placental weights and fetus-placental weight ratios were analysed as mix-model with SPSS (IBI) to take into account litter sizes.

### 3D Micro-CT imaging

Fetuses were collected at gd18.5 and processed through previously established protocols (Wong et al., 2012, 2013). Fetuses were selected at random from each litter euthanized on ice then washed and immersed in 4% paraformaldehyde 4°C for 24 hours and stored in PBS + 0.2% sodium azide. Prior to scanning, fetuses were incubated in hydrogel solution at 4°C for 3 days for monomer hybridization, degassed and heated at 37°C for 3 hours to induce hydrogel polymerisation. Hydrogel was then removed and fetuses were washed in 0.5M iodine solution, washed and mounted in 1% agarose and scanned using a *Nikon XT 225 micro-CT* (resolution varied between approximately 9-11μm). Image stacks were processed in Amira 6.3 (*FEI*) software. Brain volume was manually segmented through voxel intensity threshold selection to visualise organ boundaries. Brain volume segmentation included olfactory lobes, cortices, cerebellar and medullae (segmented at superior aspect of spinal cord). Femur length was measured directly from the tip of the greater trochanter to the articulating border of the medial condyle (ensuring the head of femur and the lateral condyle were approximately perpendicular). There was no statistical relationship between brain volume and resolution/storage length.

### LPS injections in pregnant mice

Mice were mated and the day of plug was considered gd0.5. LPS was injected intraperitoneally at gd7.5 (0.02mg/g LPS *E coli* 055:B5 (Sigma) to induce ~50% embryo death. At gd9.5 uterine tissues were collected, and embryo death, identified as black or red and dysmorphic, was evaluated. Uteri were then processed for mRNA extraction and RT-qPCR or flow cytometry. Alternatively, dams were injected with LPS at gd8.5 and uteri and blood samples were collected at gd9.5. Tissues were homogenised in 150mM NaCl, 20mM Tris, 1mM EGTA, 1% Triton X-100, 2mM PMSF, phosphatase inhibitors (Sigma), and cOmplete protease inhibitor tablets (Roche). Plasma was obtained from blood samples by centrifugation. These samples were assayed using Pro-inflammatory cytokine assay (Mesoscale) or processed for mRNA extraction and RT-qPCR or flow cytometry. Uterine tissues were dissected and processed to extract myeloid cells (as described).

### Statistical analysis

Graphs and statistics have been generated with Prism 7 (Graphpad). One way-Anova or Kruskal Wallis was performed when we compared multiple variables or RM one-way ANOVA when multiple variables were paired. Student *t*-test or Mann-Witney test was then performed when two variables were compared. Paired Student *t*-test or Wilcoxon test was performed when paired data were analysed. All data are presented as mean ± SEM and we represented p-value summary as: ns, not significant; *, P < 0.05; **, P < 0.01; ***, P<0.001; ****, P < 0.0001.

## Author Contribution

EB designed and performed experiments, analysed data, wrote the manuscript. BMJR designed and performed experiments and edited manuscript. RSH did the computational analysis of the RNAseq data and edited manuscript. NS, HEJY, JLT and IA designed and performed experiments and analysed data. ANSP designed experiments and analysed data. JK designed experiments and wrote early drafts of the manuscript. DAH, ST and SV performed experiments. FG designed and performed experiments, analysed data and edited manuscript. ANJM designed experiments and edited manuscript. FC designed experiments, analysed data and wrote manuscript.

