## Supplementary figures and images for "Maternal group 2 innate lymphoid cells contribute to fetal growth and protection from endotoxin-induced abortion in mice"

### Suppl Figures 1-7

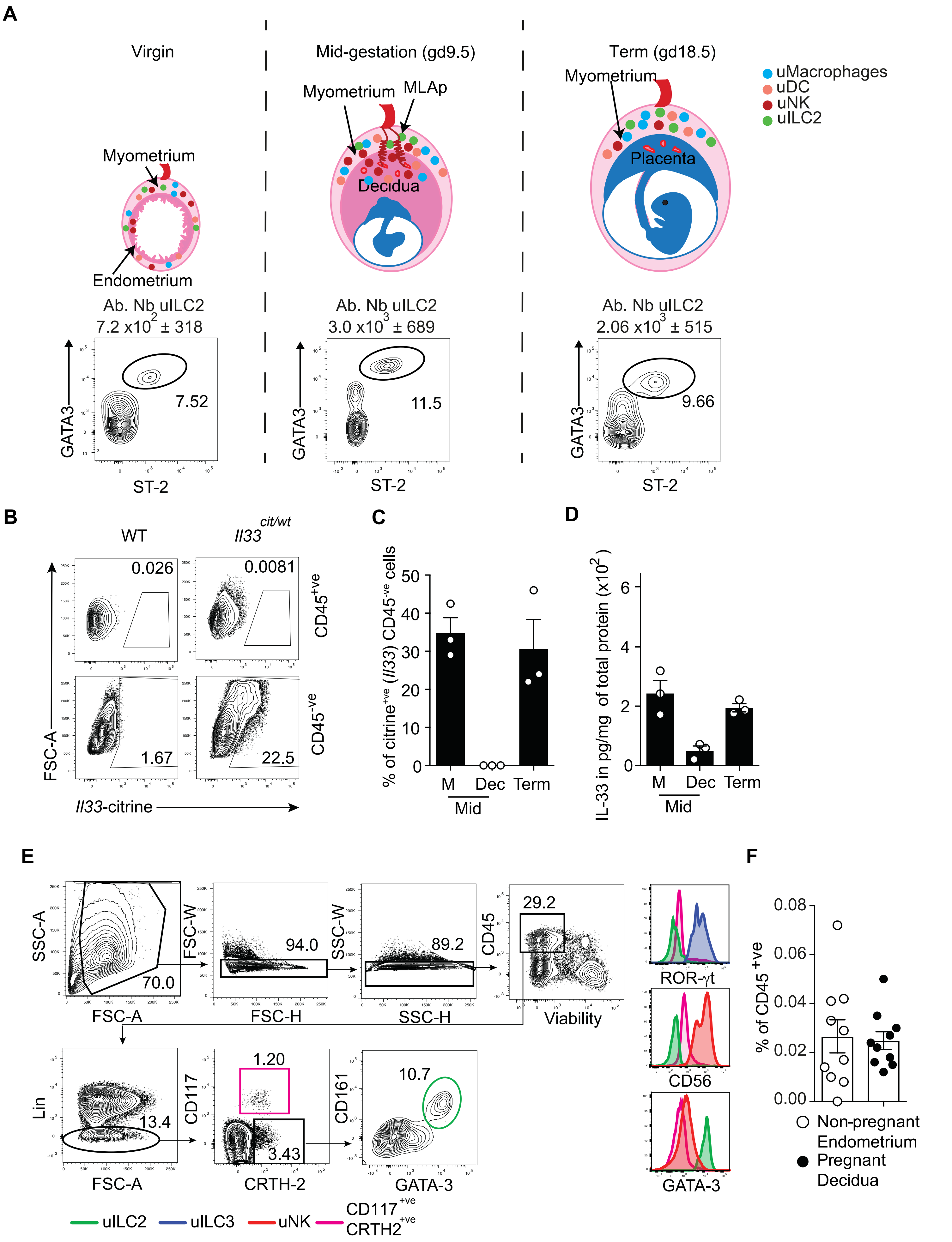

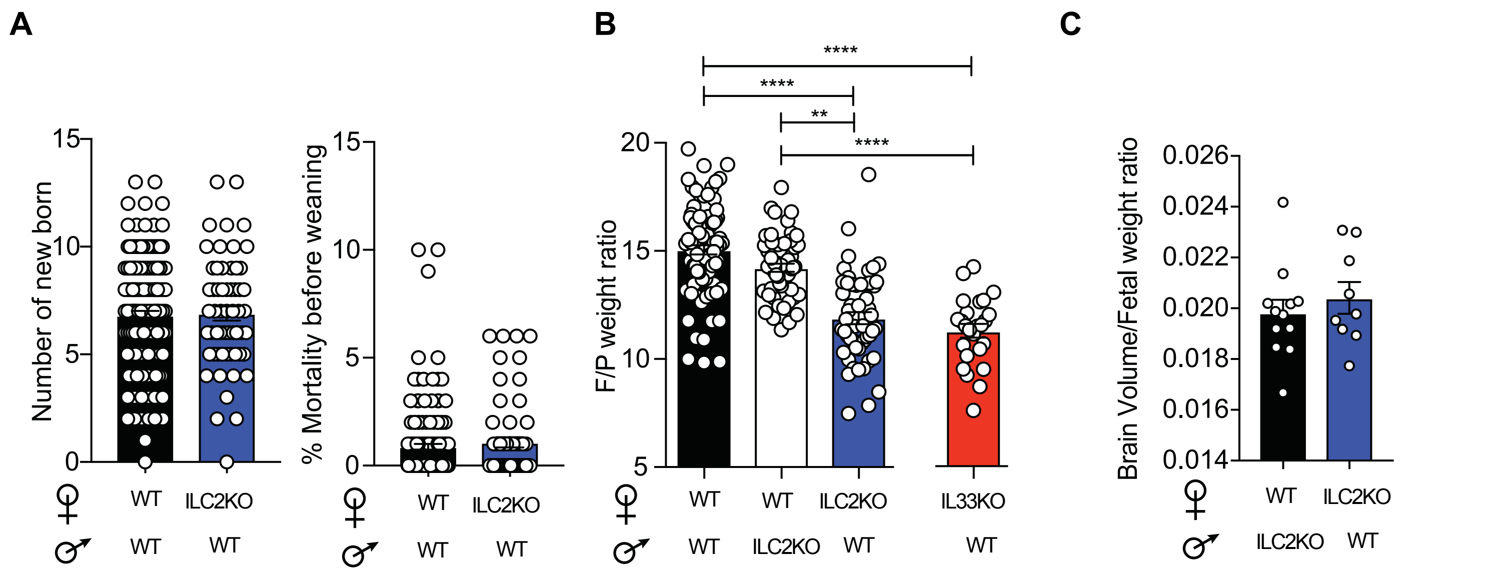

Figure S3

**A**

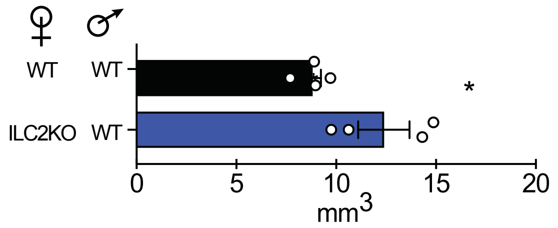

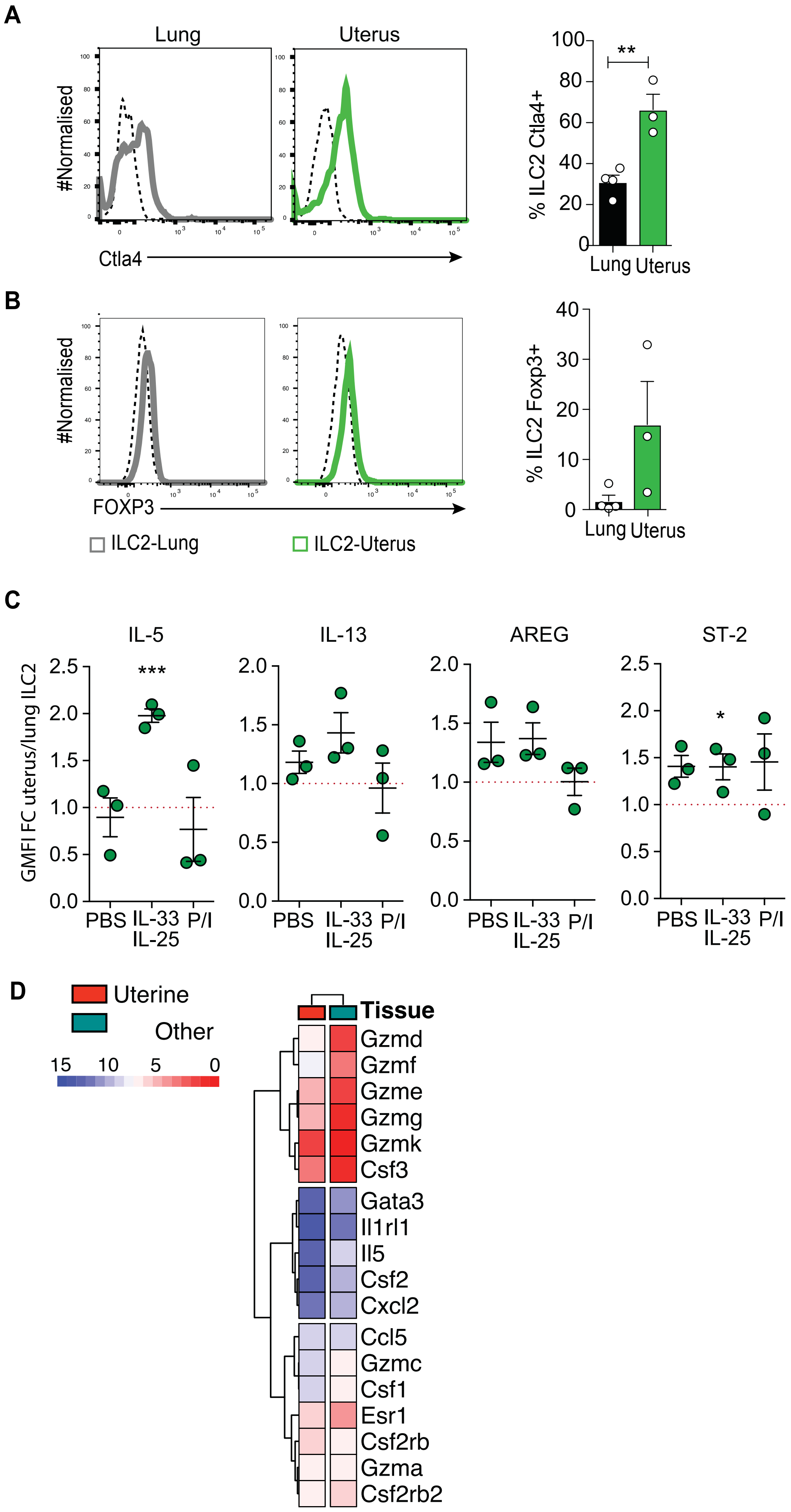

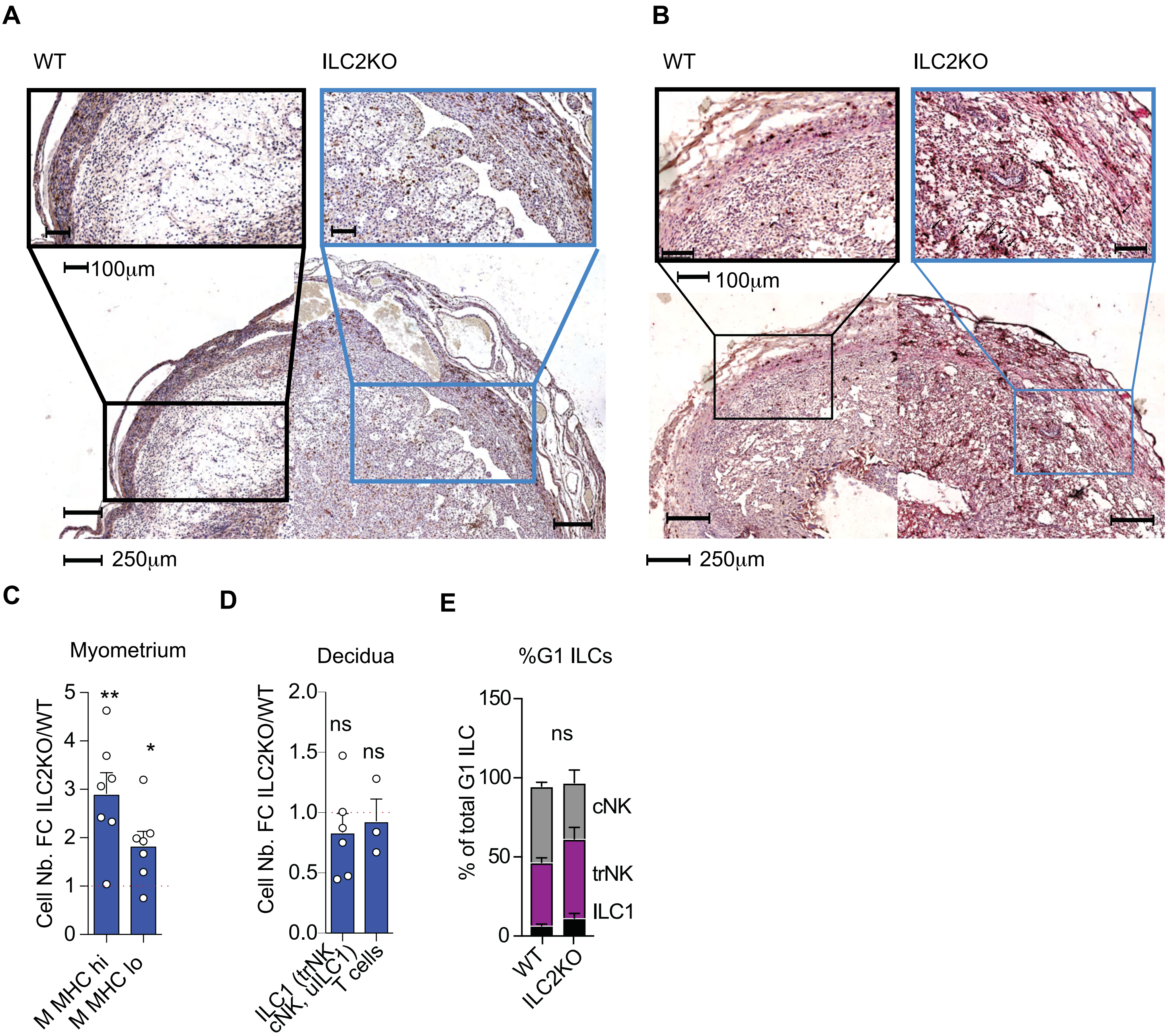

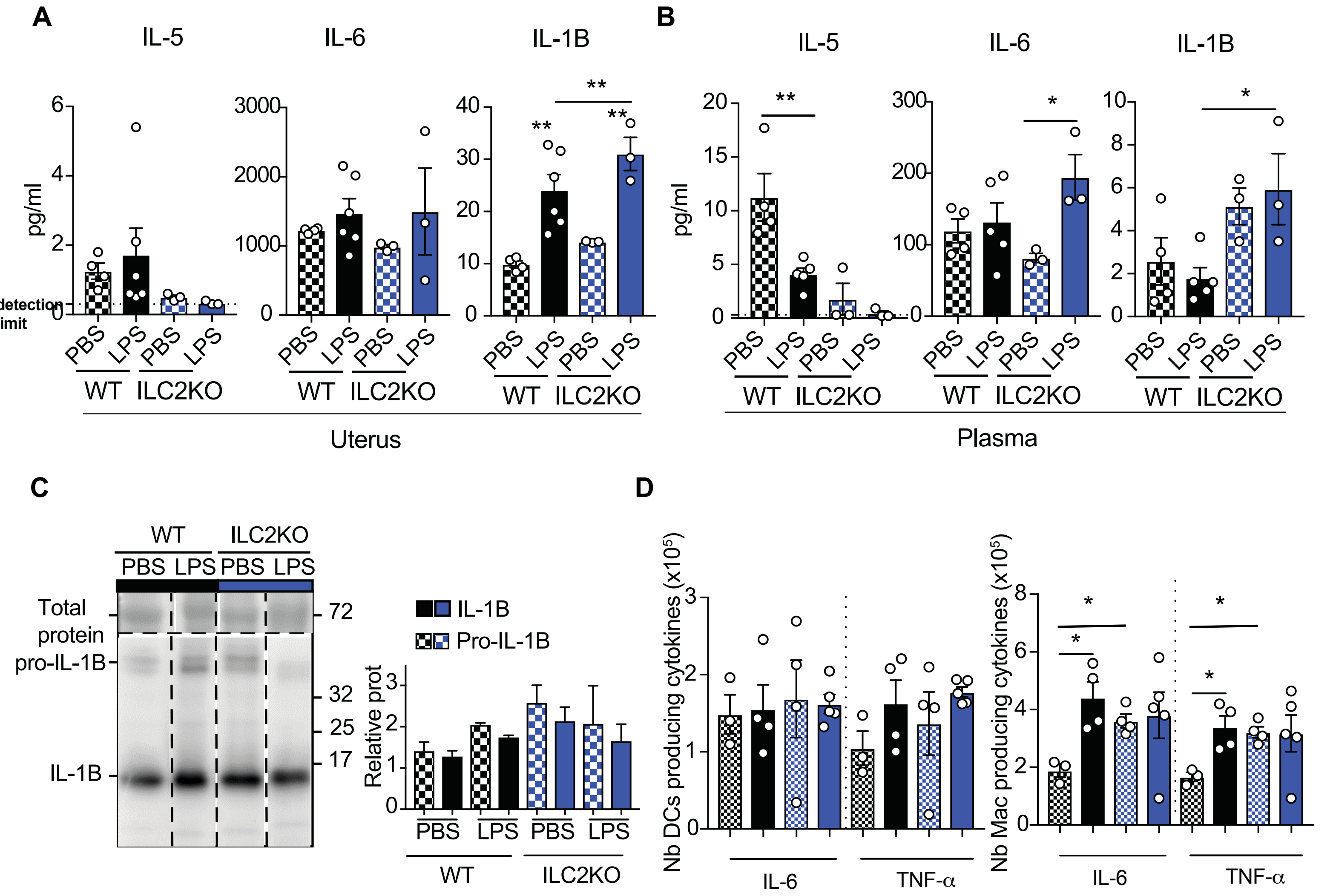

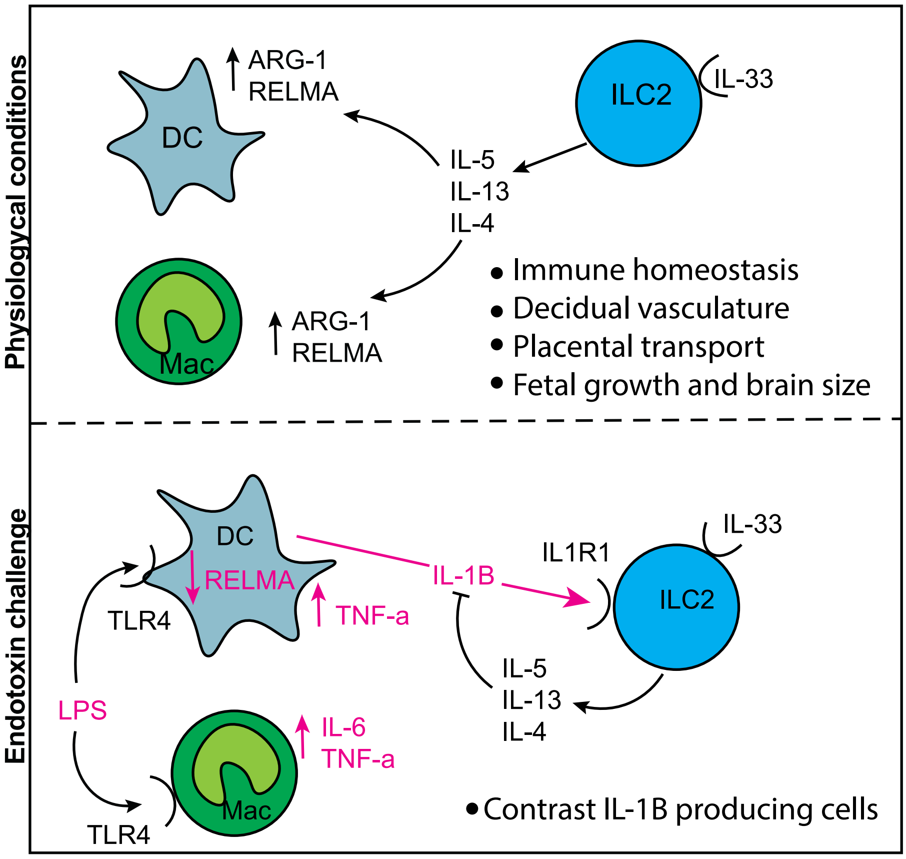
